# Studying time-resolved functional connectivity via communication theory: on the complementary nature of phase synchronization and sliding window Pearson correlation

**DOI:** 10.1101/2024.06.12.598720

**Authors:** Sir-Lord Wiafe, Nana O. Asante, Vince D. Calhoun, Ashkan Faghiri

## Abstract

Time-resolved functional network connectivity (trFNC) assesses the time-resolved coupling between brain regions using functional magnetic resonance imaging (fMRI) data. This study aims to compare two techniques used to estimate trFNC, to investigate their similarities and differences when applied to fMRI data. These techniques are the sliding window Pearson correlation (SWPC), an amplitude-based approach, and phase synchronization (PS), a phase-based technique. To accomplish our objective, we used resting-state fMRI data from the Human Connectome Project (HCP) with 827 subjects (repetition time: 0.7s) and the Function Biomedical Informatics Research Network (fBIRN) with 311 subjects (repetition time: 2s), which included 151 schizophrenia patients and 160 controls. Our simulations reveal distinct strengths in two connectivity methods: SWPC captures high-magnitude, low-frequency connectivity, while PS detects low-magnitude, high-frequency connectivity. Stronger correlations between SWPC and PS align with pronounced fMRI oscillations. For fMRI data, higher correlations between SWPC and PS occur with matched frequencies and smaller SWPC window sizes (∼30s), but larger windows (∼88s) sacrifice clinically relevant information. Both methods identify a schizophrenia-associated brain network state but show different patterns: SWPC highlights low anti-correlations between visual, subcortical, auditory, and sensory-motor networks, while PS shows reduced positive synchronization among these networks. In sum, our findings underscore the complementary nature of SWPC and PS, elucidating their respective strengths and limitations without implying the superiority of one over the other.

**Impact Statement:** This study demonstrates that SWPC and PS provide complementary insights into dynamic functional connectivity, revealing different aspects of brain dynamics based on signal focus. For tasks involving slow dynamics, SWPC amplitude is ideal, while the PS phase is more suitable for transient dynamics. In schizophrenia, typically associated with general dysconnectivity, we uncover a dual dysconnectivity profile depending on phase or amplitude dynamics. This novel approach offers researchers a platform to explore task-specific dysconnectivity profiles, enabling more targeted interventions. These findings will guide methodology choices, deepen understanding of brain dynamics, and support the development of precise neuropsychiatric biomarkers.

**Highlights:**

- Time-resolved functional network connectivity (trFNC) is widely used; here we study two approaches often pit against one another: 1) phase synchrony (PS), a phase-based method, and 2) sliding window Pearson correlation (SWPC), an amplitude-based method.
- SWPC is sensitive to the choice of window size, while PS requires a narrow frequency band. Both can result in the loss of relevant information.
- We find through simulation that SWPC better captures high-magnitude slow-varying amplitude-encoded connectivity while PS better captures low-magnitude fast-varying phase-encoded connectivity.
- We find that while both SWPC and PS detect disconnected states mostly associated with schizophrenia they exhibit unique complementary patterns.
- We conclude that SWPC and PS are complementary techniques, each with distinct assumptions and constraints, which should be selected based on the focus of the study.

## 1. INTRODUCTION

The study of functional connectivity (FC) offers significant insights into inter-network communication within the brain using functional magnetic resonance imaging (fMRI) (Biswal, Yetkin et al. 1995, Lowe, Dzemidzic et al. 2000, Fingelkurts, Fingelkurts and Kähkönen 2005, Bastos and Schoffelen 2016). FC and functional network connectivity (FNC) - functional connectivity between whole brain intrinsic fMRI networks have uncovered several key findings about intrinsic blood oxygenation level-dependent (BOLD) brain activity, particularly through resting-state fMRI (rsfMRI), and have provided valuable information about various brain disorders, including but not limited to schizophrenia (Calhoun, Kiehl et al. 2004, Lynall, Bassett et al. 2010, Sheffield and Barch 2016), Alzheimer’s disease (Allen, Barnard et al. 2007, Chhatwal, Schultz et al. 2013, Sheline and Raichle 2013), and depression (Veer, Beckmann et al. 2010, Zeng, Shen et al. 2012, Mulders, van Eijndhoven et al. 2015). Numerous methods exist for quantifying FC/FNC between brain network time series, including Pearson correlation (Greicius, Krasnow et al. 2003, Damoiseaux, Rombouts et al. 2006), partial correlation (Sun, Miller and D’esposito 2004), mutual information (Calhoun, Kim and Pearlson 2003, Wang, Alahmadi et al. 2015, Salman, Vergara et al. 2019, Mohanty, Sethares et al. 2020, Motlaghian, Belger et al. 2022), phase locking value (Zhang, Pan et al. 2016), and dynamic time warping (Wiafe, Faghiri et al. 2024, Wiafe, Faghiri et al. 2024, Wiafe, Kinsey et al. 2024, Wiafe, Kinsey et al. 2025), among others.

Over the past few years, there has been a growing body of research extending FC studies to capture FC in a temporally resolved manner (Chang and Glover 2010, Sakoğlu, Pearlson et al. 2010, Hutchison, Womelsdorf et al. 2013, Allen, Damaraju et al. 2014). This approach, often referred to as dynamic functional connectivity (Calhoun, Miller et al. 2014, Iraji, Faghiri et al. 2020) (Hutchison, Womelsdorf et al. 2013, Lurie, Kessler et al. 2020), reveals how brain networks are coupled over time. Supported by the notion that the brain is a highly complex dynamic system (Kelso 1995, Telesford, Simpson et al. 2011) with intricate inter-network connections (Achard, Salvador et al. 2006, Van den Heuvel and Sporns 2013, Sporns 2016) across different spatiotemporal scales (Majeed, Magnuson and Keilholz 2009, Iraji, Fu et al. 2019), trFNC has provided a deeper understanding of brain function and organization (Hutchison, Womelsdorf et al. 2013, Allen, Damaraju et al. 2014).

Several methods for estimating trFNC exist, such as sliding window Pearson correlation (SWPC) (Allen, Damaraju et al. 2014, Leonardi and Van De Ville 2015), cross wavelet coherence (Chang and Glover 2010, Yaesoubi, Allen et al. 2015), phase synchrony (PS) (Pedersen, Omidvarnia et al. 2018, Honari, Choe et al. 2020), coactivation patterns (Eickhoff, Bzdok et al. 2011, Liu, Zhang et al. 2018), hidden Markov models (Zhang, Cai et al. 2020, Hussain, Langley et al. 2023) and other window-based approaches based on Spearman and Kendall correlation (Savva, Mitsis and Matsopoulos 2019, Torabi, Mitsis and Poline 2024), and mutual information (Savva, Mitsis and Matsopoulos 2019, Torabi, Mitsis and Poline 2024),

Traditionally recognized as a prominent measure of time-resolved functional connectivity in the field of neuroscience, SWPC excels in quantifying the time-varying connectivity of different brain regions (Sakoğlu, Pearlson et al. 2010, Handwerker, Roopchansingh et al. 2012, Jones, Vemuri et al. 2012, Tagliazucchi, Von Wegner et al. 2012, Hutchison, Womelsdorf et al. 2013, Hutchison, Womelsdorf et al. 2013, Keilholz, Magnuson et al. 2013, Allen, Damaraju et al. 2014, Calhoun, Miller et al. 2014, Zalesky, Fornito et al. 2014, Du, Fryer et al. 2018, Faghiri, Stephen et al. 2018, Mokhtari, Akhlaghi et al. 2019, Iraji, Faghiri et al. 2020, Faghiri, Adali and Calhoun 2022, Kazemivash and Calhoun 2022, Faghiri, Yang et al. 2023). While SWPC has gained considerable acceptance as a reliable measure for brain connectivity, it is important to note that it specifically captures linear co-variations that are related to the signal. This means that SWPC is adept at identifying instances where brain regions activate simultaneously in a linear and directly proportional manner.

Recently, Pedersen et al. compared SWPC with a more recently used technique for fMRI data, phase synchrony (Pedersen, Omidvarnia et al. 2018). PS measures the synchronization of neural activities based on the timing of their phase relationships (Lachaux, Rodriguez et al. 1999, Mormann, Lehnertz et al. 2000, Laird, Carew and Meyerand 2001, Varela, Lachaux et al. 2001, Laird, Rogers et al. 2002, Deshmukh, Shivhare et al. 2004, Costa, Rognoni and Galati 2006, Pockett, Bold and Freeman 2009, Fell and Axmacher 2011, Glerean, Salmi et al. 2012, Sun, Hong and Tong 2012, Bolt, Nomi et al. 2018, Kumar, Reddy and Behera 2018, Pedersen, Omidvarnia et al. 2018, Honari, Choe et al. 2020). PS is typically computed by estimating the phase of the signal and finding the phase difference. The most commonly used approach to estimating the instantaneous phase of a signal is to employ the analytic signal approach based on the Hilbert transform (Bedrosian 1962) while others use wavelet transforms (Boashash 1992).

Pedersen et al. compared these two methods by quantifying the average Spearman correlation between PS and SWPC with varying window sizes, highlighting their similarities using fMRI data (Pedersen, Omidvarnia et al. 2018). However, the PS method they used employed an all-positive phase synchronization measure, overlooking anti-synchronization estimations. To make PS comparable to SWPC, Pedersen et al. computed the correlation between the all-positive PS measure and the absolute values of SWPC (Pedersen, Omidvarnia et al. 2018). Honari et al. emphasized the benefits of using a PS measure that accounts for both positive and negative values by employing the cosine function instead of the sine function (Honari, Choe et al. 2020). Additionally, while Pedersen’s findings are insightful, averaging the correlations between the two methods does not provide insights into their temporal dependencies. In magnetoencephalography (MEG) studies, the concepts of amplitude coupling and phase coupling have been shown to reflect at least partially distinct neuronal mechanisms (Siems and Siegel 2020). Furthermore, our previous study shows that the correlation between SWPC and PS is time-dependent (Wiafe, Fu et al. 2023), indicating the need for a more nuanced analysis to understand the time-resolved relationships between these measures.

Our primary objective is to compare these two methods in a time-varying manner while addressing the above concerns. In contrast to earlier work claiming that phase synchrony is preferable to SWPC (Pedersen, Omidvarnia et al. 2018), we show through simulations and fMRI analysis that these two methods offer complementary insights into time-resolved functional connectivity. Also, from the perspective of signals and communication theory, these methods depend on different fundamental aspects of a signal: SWPC is amplitude-based, while PS is phase-based. By studying trFNC through the lens of communication theory, we aim to shed more light on the complementary information offered by these two methods. This approach can provide deeper insights into how brain networks communicate and interact over time, shedding light on their time-resolved relationships and functional organization. Our study employs independent component analysis (ICA) to extract intrinsic brain networks from the fMRI data (Calhoun, Adali et al. 2001, Du, Fu et al. 2020).

Our paper is structured as follows: a background section drawing parallels between SWPC and PS with communication system demodulation; simulations illustrating the complementary insights each method provides; an analysis of time-resolved correlations between SWPC and PS in relation to the spectral characteristics of fMRI data across two datasets; clinically relevant findings for Schizophrenia; and a discussion emphasizing the complementary, non-hierarchical nature of the two approaches. MATLAB code for all analyses is provided.

## 2. BACKGROUND

Some studies have noted similarities between trFNC measures: SWPC with amplitude demodulation (Faghiri, Iraji et al. 2022) and PS with phase demodulation (Wang and Da 2012). In communication theory, modulation is a technique used to transmit signals effectively from one point to another (Haykin and Moher 1989). In communication theory, modulation transmits signals by altering a carrier wave based on a message signal (Haykin and Moher 1989, Roden 1991). Analog modulation schemes such as amplitude (AM), frequency (FM), and phase (PM) modulation vary the carrier’s amplitude, frequency, or phase, respectively, to embed information (Smillie 1999, Sharma, Mishra and Saxena 2010). The canonical carrier, *c(t)*, and message signal, *m(t)*, are

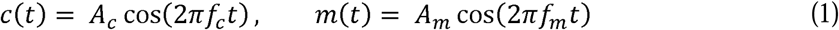

Where, *A_C_, A_m_* and *r_C_, r_m_* denote the amplitudes and frequencies of carrier and message, respectively.

In AM the composite waveform

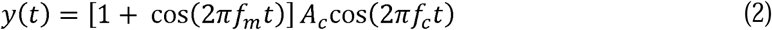

is a linear modulation scheme that follows the superposition principle. When multiple signals modulate a carrier, the resulting AM signal equals the sum of the individual AM signals (Haykin and Moher 1989, Proakis, Salehi et al. 1994).

SWPC and amplitude demodulation share similar subsystems (Faghiri, Iraji et al. 2022). Due to this similarity, using a time-varying method like the sliding window Pearson correlation (SWPC) is effective for analyzing signals in line with the time-varying nature of the message signal *m(t).* In this method, high correlation between the carrier *c(t),* and the modulated signal *y(t)* indicates high message signal amplitude, while low correlation suggests lower amplitude. This demonstrates the usefulness of correlation in AM demodulation. Given the carrier signal, *c(t) A A_C_cos2*π*r_C_ t*, and the modulated signal, *y(t) A* 11 + 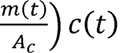, the SWPC between *y(t)* and *c(t)* approximates the amplitude variation of *m(t):*

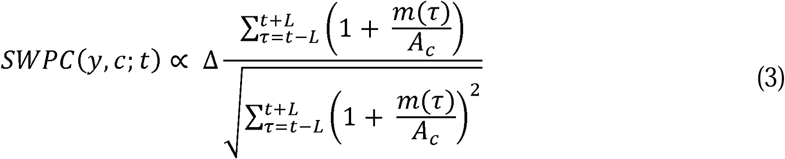

Thus, SWPC approximates a normalized form of the message signal by quantifying how closely the modulated signal aligns with the carrier over time (see supplementary material 1 for full proof).

Phase modulation (PM) encodes the message signal’s amplitude into the phase of the carrier. Due to the nonlinear nature of the cosine function, PM is inherently nonlinear (Haykin and Moher 1989, Proakis, Salehi et al. 1994). In PM, information is carried in the signal’s phase and recovered through phase demodulation. Phase synchrony methods, which compare the instantaneous phases of two signals, align with the logic of phase demodulation. For a modulated signal *y(t)*, its instantaneous phase is 2*πr_C_ t*+ *m(t)*. By subtracting the phase of the unmodulated carrier, 2*πr_C_ t*, the original message signal can be recovered:

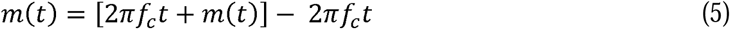

This approach depends on precise phase difference measurements. However, due to phase wrapping at 2π, small changes near these points can lead to discontinuities, introducing nonlinearities that are not present in amplitude-based methods.

SWPC and phase synchrony are distinct methods for analyzing neural activity. SWPC focuses on signal strength relationships, revealing co-activation patterns across brain regions, while phase synchrony examines timing and phase alignment, highlighting neural synchronization. We hypothesize that comparing these methods will illuminate the diverse neural mechanisms underlying cognition. Different cognitive tasks may involve precise timing without concurrent high activation, or vice versa. Recognizing such patterns could enhance our understanding of how the brain mobilizes networks for different functions.

## 3. DATA & DATA PROCESSING

### 3.1. fMRI data

This study received approval from an ethics board, and all participants provided consent by signing a form approved by the institutional review board (IRB).

The first dataset used is the Function Biomedical Informatics Research Network (fBIRN) study, which includes resting fMRI data collected from schizophrenia patients and controls. Scans were collected at a repetition time (TR) of 2 seconds. The study led to a total of 160 controls with an average age of 37.04 ± 10.86 years, ranging from 19 to 59 years. Among these, 45 were female and 115 were male. Additionally, there were 151 patients diagnosed with schizophrenia, with an average age of 38.77 ± 11.63 years, ranging from 18 to 62 years. In this group, 36 were female and 115 were male. The typical controls and schizophrenic patients were meticulously matched in terms of age, gender distribution, and mean framewise displacement during scans (age: p = 0.18; gender: p = 0.39; mean framewise displacement: p = 0.97). Schizophrenic patients were in a clinically stable condition during the time of their scans. The fBIRN dataset is used to assess the similarity between SWPC and PS in relation to its clinical relevance to the brain disorder, schizophrenia.

The second dataset used in our study is the resting-state fMRI dataset collected from 827 subjects via the Human Connectome Project (HCP) database (Van Essen, Ugurbil et al. 2012, Van Essen, Smith et al. 2013). We utilized the first-session scans acquired using a Siemens Skyra 3T scanner with a multiband accelerated, gradient-echo echo-planar imaging (EPI) sequence. The scanning parameters were set to a repetition time (TR) of 0.72s, 72 slices, an echo time (TE) of 58ms, and a flip angle of 90°. A voxel size of 2 × 2 × 2 mm was used to acquire 1200 time points of fMRI data for each subject. Given the HCP dataset has an extensive participant pool, we use it as a validation dataset for the similarity between SWPC and PS.

### 3.2. fMRI pre-processing

fMRI data require extensive preprocessing to correct for various sources of noise and artifacts before analysis. Data was preprocessed using <u>statistical parametric mapping</u> 5 (SPM; http://www.fil.ion.ucl.ac.uk/spm (Friston 2003). These preprocessing steps commonly include slice timing correction, realignment, spatial normalization, and spatial smoothing (Turner, Howseman et al. 1998, Penny, Friston et al. 2011, Esteban, Markiewicz et al. 2019). Following preprocessing (Fu, Batta et al. 2024) we implemented the NeuroMark pipeline, a fully automated spatially constrained ICA on the preprocessed fMRI data (Du, Fu et al. 2020). Using the neuromark_fMRI_1.0 template (available in GIFT at http://trendscenter.org/software/gift or at http://trendscenter.org/data) we generated 53 intrinsic connectivity networks (ICNs) for each subject. These ICNs are grouped into brain functional domains including Subcortical (SC), Auditory (Aud), Sensorimotor (SM), Visual (Vis), Cognitive Control (CC), Default Mode (DM), and Cerebellum (Cb).

### 3.3. fMRI post-processing

To enhance the quality of these ICNs, we implemented detrending and despiking methods to eliminate drifts, sudden fluctuations, and significant artifacts that may have persisted after the initial preprocessing stages. The time series of the ICNs were bandpass-filtered in the range of 0.01 to 0.15 Hz, a frequency range in fMRI research that is suggested to be relevant for identifying brain domain BOLD signals (Braun, Plichta et al. 2012, Yaesoubi, Allen et al. 2015). An infinite impulse response (IIR) filter was designed using the butter function in MATLAB and applied via the filtfilt function to ensure zero phase shifts and preserve phase information (Oppenheim 1999, Proakis 2007, Caballero-Gaudes and Reynolds 2017). The optimal filter order was estimated using the buttord function in MATLAB, which returns the lowest order of the Butterworth filter with no more than 3dB ripple in the passband and at least 30dB of attenuation

in the stopband. Finally, we performed z-scoring on the ICNs. The MATLAB version used was MATLAB R2022a for all steps of the analysis.

## 4. METHOD

### 4.1. Sliding window Pearson correlation

As previously discussed in the background section, the SWPC method calculates the Pearson correlation between signal pairs, sliding a defined window by one time point until covering all time points. This method is a standard approach for computing trFNC between brain regions (Hutchison, Womelsdorf et al. 2013, Allen, Damaraju et al. 2014, Mokhtari, Akhlaghi et al. 2019, Iraji, Faghiri et al. 2020, Faghiri, Iraji et al. 2022, Wiafe, Fu et al. 2023). Given a time series pair, *x(t)* and *y(t)* the sliding window Pearson correlation over a defined window of length, 2L + 1, is given below:

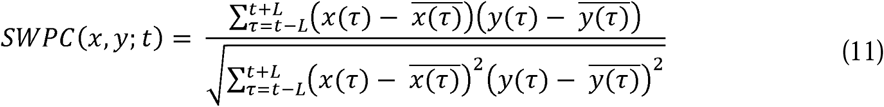

A key challenge in using SWPC is determining the optimal window length to effectively capture connectivity between a signal pair. SWPC’s sensitivity to window length has been well-documented (Hutchison, Womelsdorf et al. 2013). The selected window must be large enough to detect low connectivity yet small enough to identify transient connectivity (Sakoğlu, Pearlson et al. 2010, Leonardi and Van De Ville 2015). Leonardi and Van De Ville recommend using a minimum window length longer than the longest wavelength of the signal, approximately 100 seconds, particularly when the low-frequency cutoff is 0.01 Hz. Other studies have suggested that shorter window lengths, ranging from 20 to 60 seconds, could be viable for estimating trFNC in fMRI data (Shirer, Ryali et al. 2011, Zalesky and Breakspear 2015, Liégeois, Ziegler et al. 2016, Preti, Bolton and Van De Ville 2017, Pedersen, Omidvarnia et al. 2018).

In our study, we opt for a window size aligned with the -3dB cutoff frequency of the high-pass rectangular window filter subsystem of SWPC (Faghiri, Iraji et al. 2021, Faghiri, Yang et al. 2023, Wiafe, Fu et al. 2023), as detailed below:

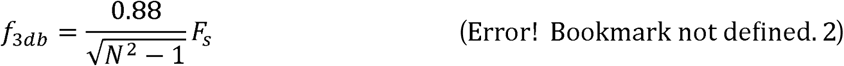

where N is the window size and *F_S_* is the sampling frequency (1/0.72s =1.39hz for HCP and 1/2s = 0.5hz for fBIRN). Due to the application of a bandpass filter on the data, the high-pass cutoff *f*_3*db*_ should be higher than 0.01Hz, which results in a minimum N value of ∼123 samples (∼88s) for HCP and ∼44 samples (∼88s) for fBIRN.

### 4.2. Phase Synchrony

Phase synchrony or synchronization is a phenomenon where the phases of two oscillating signals align over time. This concept has been instrumental in exploring functional connectivity, as evidenced by various studies (Lachaux, Rodriguez et al. 1999, Fell and Axmacher 2011, Glerean, Salmi et al. 2012, Omidvarnia, Pedersen et al. 2016, Bolt, Nomi et al. 2018, Pedersen, Omidvarnia et al. 2018). There are two primary approaches for analyzing phase synchrony: one involves a window-based approach, like SWPC, while the second method involves using the estimated instantaneous phase of signal pairs. The window-based approaches include techniques like phase locking value (Tass, Rosenblum et al. 1998, Rebollo, Devauchelle et al. 2018), circular-circular correlation (Jammalamadaka, Sengupta and Sengupta 2001), and Toroidal-Circular Correlation (Sojakova 2016, Zhan, Ma et al. 2019). The instantaneous phase approach employs metrics such as phase coherence, where the sine of the phase difference is employed (Pedersen, Omidvarnia et al. 2017, Pedersen, Omidvarnia et al. 2018) and the cosine of relative phase, where the cosine of phase difference is employed (Cabral, Vidaurre et al. 2017, Honari, Choe et al. 2020, Vohryzek, Deco et al. 2020, Hancock, Cabral et al. 2022, Wiafe, Fu et al. 2023). Honari et al. provide comprehensive details on these methods and approaches.

In our study, we adopt the instantaneous phase approach, which necessitates estimating the instantaneous phase of signals for analyzing phase synchrony. We specifically utilize the cosine of the phase difference because it provides a readily comparable measure to SWPC. Positive values indicate high synchronization, while negative values suggest anti-synchronization, akin to correlation coefficients. Given a narrow-banded time series pair, *x(t)* and *y(t),* phase synchrony computed by the cosine of relative phase is given below:

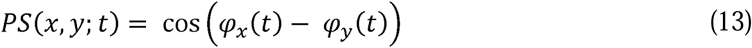

Where φ*_X_ (t) A arctan(x(t) + JH x(t)J)* and *H ·J* represents the Hilbert transform. The Hilbert transform is commonly used for this estimation, provided that the Bedrosian theorem’s conditions are met. According to the Bedrosian theorem, successful phase extraction requires the spectral properties of the signal’s envelope and phase to be separate and distinct from each other (Bedrosian 1962, Xu and Yan 2006). The narrower the bandwidth of the signal, the more likely the Hilbert transform accurately models the signal (Boashash 1992, Delprat, Escudié et al. 1992, Picinbono 1997, Chavez, Besserve et al. 2006).

In this study, where the primary focus is to compare the performance of SWPC and phase synchrony, we adhere to standard practices concerning the choice of frequency band and bandwidth in fMRI analysis in phase synchrony measures. Extensive research suggests that the frequency band [0.04, 0.07JHz is commonly regarded as the most reliable and functionally relevant for capturing meaningful neural activity (Biswal, Yetkin et al. 1995, Achard, Salvador et al. 2006, Glerean, Salmi et al. 2012). Glerean, Salmi et al. specifically employed this frequency band for computing phase synchrony due to its characteristics of yielding robust fMRI signals least affected by noise, as supported by prior studies (Biswal, Yetkin et al. 1995, Achard, Salvador et al. 2006, Buckner, Sepulcre et al. 2009). Furthermore, some studies have utilized a slightly broader frequency band of [0.03, 0.07]*Hz* (Pedersen, Omidvarnia et al. 2017, Pedersen, Omidvarnia et al. 2018). Pedersen, Omidvarnia et al. compared SWPC to phase synchrony using a measure based on 1 - ahs(sin (*φ_x_X(t) - q,y(t*))), emphasizing the strong relationship between SWPC and phase synchrony within the frequency band [0.03, 0.07]*Hz*. We opt for the same frequency band [0.03, 0.07JHz to highlight both the disparities and complementary aspects between SWPC and phase synchrony methods rather than focusing on their similarities, as demonstrated by Pedersen, Omidvarnia et al., given our study’s alignment with their objectives.

### 4.3. SWPC and PS using synthesized data

We assessed SWPC and phase synchrony using synthesized data across three scenarios to highlight their complementary nature. In the first scenario, we show that phase synchrony effectively captures connectivity characterized by weak coupling at high frequencies, where SWPC may not perform as well. Phase-based methods excel in examining synchronization among weakly coupled oscillators, supported by studies like the Kuramoto model and others (Rosenblum, Pikovsky and Kurths 1996, Kopell and Ermentrout 2002, Dorfler and Bullo 2012, Schwemmer and Lewis 2012, Thümler, Srinivas et al. 2023). This aligns with findings that weak coupling is best explained by “phase-only” models (Fagerholm, Moran et al. 2020) and that weak synaptic coupling can show high-frequency synchronization (Lee and Sepulchre 2024). We generated time-resolved connectivity ground truth with low magnitudes and high frequencies to demonstrate phase synchrony’s superiority over SWPC.

In the second scenario, we show that SWPC captures low-frequency connectivity with strong coupling, which phase synchrony may miss. Fagerholm et al. found that anesthetized brain states depend on oscillator amplitudes, favoring “amplitude-only” models (Fagerholm, Moran et al. 2020). This aligns with studies showing simpler, more stable anesthetic neuronal dynamics (Scott, Fagerholm et al. 2014, Fagerholm, Scott et al. 2016). They also demonstrated that phase models fail with strong coupling (Fagerholm, Moran et al. 2018). Thus, we generated connectivity ground truth with high magnitudes and low frequencies to highlight SWPC’s advantages.

In the third scenario, we aimed to merge scenarios 1 and 2 by creating time-resolved connectivity characterized by mid-strength and mid-frequency range. Both phase synchrony and SWPC are anticipated to accurately estimate connectivity in this scenario.

#### 4.3.1. Scenario 1: Weak high-frequency connectivity encoded in phase

We initiate the procedure by generating two bandlimited signals, *x(t)* and *y(t). x(t)* and *y(t*) can be expressed as below:

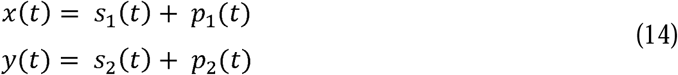

For *p_l_(t),* we generate a high-frequency bandlimited phase signal in the frequency range of [0.05, 0.15]*Hz* from a Gaussian distribution with a mean of zero, expressed as *p_l_(t) A sin(2πrt + q,(t) + c(t)).* Here r denotes the central frequency, *q,(t)* represents a random phase deviation signal, and *c(t)* is the connectivity signal which is a high-frequency bandlimited signal with frequency range of [0.02, 0.03]*Hz* with a maximum amplitude of 0.3 also generated from a Gaussian distribution with a mean of zero. We encode the connectivity into the phase of *p_l_(t),* by combining *c(t)* with *φ(t)* within *p_l_(t).* Concurrently, we generate a low-frequency random signal *s_l_(t),* bandlimited to the frequency range of [0.01, 0.03]*Hz* so that its frequency does not overlap with *p_l_(t).* The signal *y(t)* is created similarly, where we generate another low-frequency random signal *s_l_(t)* and a high-frequency phase signal *p*_2_(*t*)*A cos*(2π*ft* + *φ(t))*. We use a cosine function for *p*_2_(*t*) to ensure that the phase synchrony using the cosine of the phase difference yields *c(t).* This scenario creates a phase modulation scenario

where the message signal (time-resolved connectivity) is encoded in the phase of the signal. The process of generating scenario 1 is illustrated in Figure 1. Finally, Gaussian noise is added to both *x(t)* and *y(t)* to achieve a signal-to-noise ratio (SNR) of 20dB, which is within the reported SNR range of fMRI time series (Welvaert and Rosseel 2013). To assess SWPC and phase synchrony in this scenario, we compute SWPC across multiple window sizes and phase synchrony across multiple filtering frequency bands between 100 randomized iterations of *x(t)* and *y(t)* and compare their respective results to the ground truth connectivity *c(t)*.

**Figure 1.**
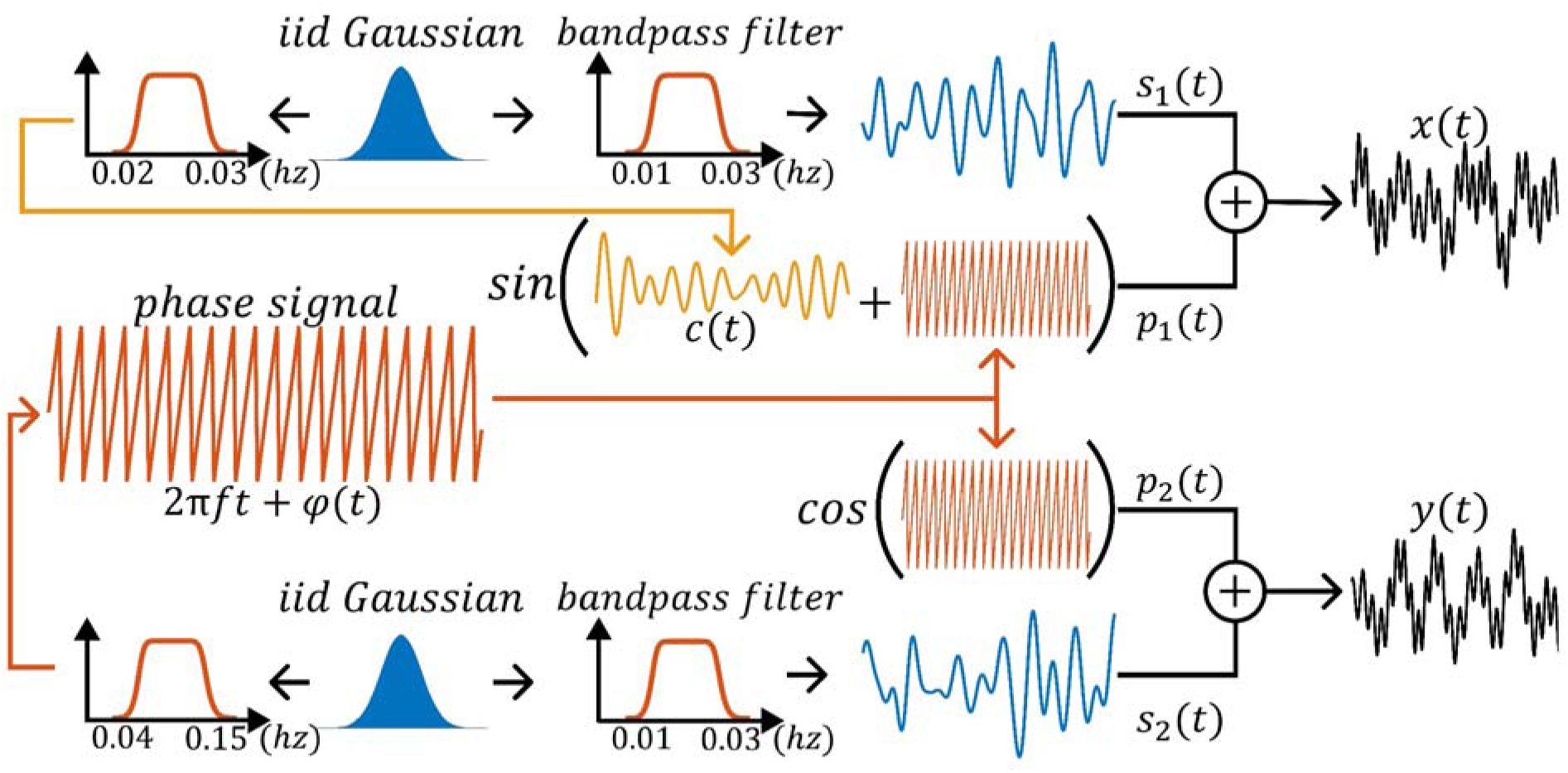
illustrates the pipeline for Scenario 1. A weak high-frequency time-resolved connectivity is encoded in the phase of the output signals. To prevent the amplitude of the signals from encoding the low-amplitude high-frequency time-resolved connectivity ground truth, random Gaussian signals with non-overlapping frequencies are introduced and added to the phase signals..

#### 4.3.2. Scenario 2: Strong low-frequency connectivity encoded in amplitude

In scenario 2, we adhere to the methodology utilized by Faghiri, Yang et al. for generating time-resolved connectivity between pairs of band-limited signals encoded in their amplitudes using Cholesky decomposition (Schmittmann, Jahfari et al. 2015, Kang, Ombao et al. 2017, Pedersen, Omidvarnia et al. 2018, Faghiri, Yang et al. 2023).This methodology involves the creation of two random time series, which are subsequently bandlimited to a frequency range of [0.01, 0.15]*Hz*. To establish a time-resolved connectivity in the amplitudes of *x(t)* and *y(t),* we generate *c(t)* as a low-frequency time-resolved connectivity signal, bandlimited to the frequency range of [0.002, 0.004]*Hz*, with an amplitude of 0.9. Subsequently, *c(t)* is utilized to construct a time-varying covariance matrix in the following form:

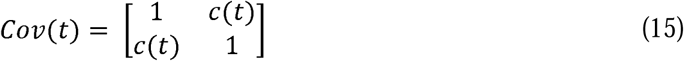

Cholesky decomposition is employed to decompose this covariance matrix at each time point. The resulting upper triangular matrix is then multiplied by *x(t)* and *y(t),* thereby encoding a time-resolved connectivity of *c(t)* at time *t* between *x(t)* and *y(t).* This scenario creates an amplitude modulation scenario where the message signal (connectivity) is encoded in the amplitude of a signal. The process of scenario 2 is illustrated in Figure 2. Similarly to scenario 1, Gaussian noise with a mean of zero and a standard deviation corresponding to an SNR of 50dB is added to *x(t)* and *y(t).* We evaluate SWPC and phase synchrony by computing SWPC across multiple window sizes and phase synchrony across multiple bandwidths between 100-sample pairs of *x(t)* and *y(t)* and comparing their respective results to the ground truth connectivity *c(t)*.

**Figure 2.**
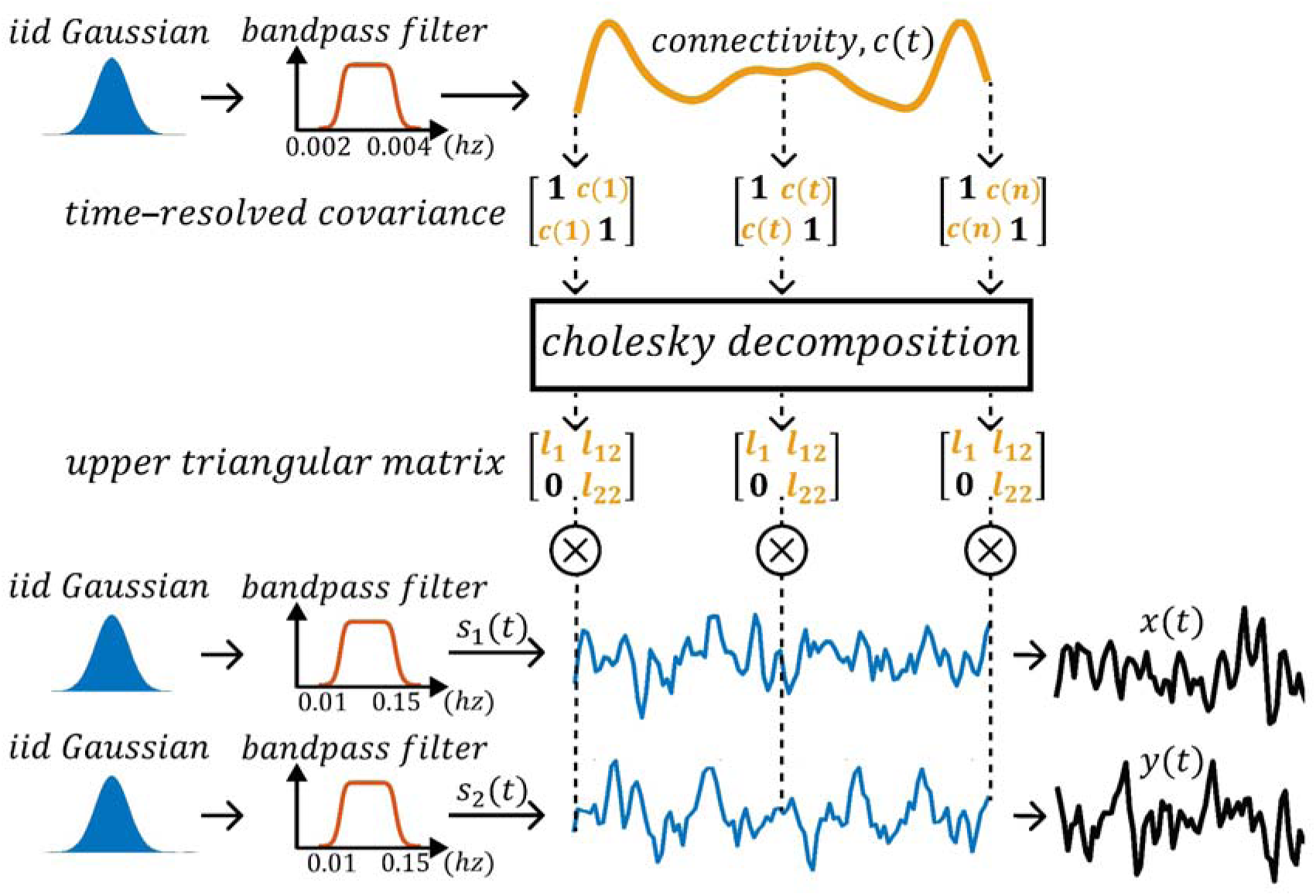
illustrates the pipeline for Scenario 2. Strong low-frequency time-resolved connectivity, modulated by the amplitude of the signals using Cholesky decomposition, is depicted. Firstly, two bandlimited random signals are generated and modulated by the time-resolved upper triangular Cholesky decomposed matrices derived from the strong low-frequency time-resolved connectivity. This process generates two signals with the ground truth time-resolved connectivity encoded in their amplitudes.

#### 4.3.3. Scenario 3: Mid-strength mid-frequency connectivity encoded in both amplitude and phase

In scenario 3, we follow a methodology similar to scenario 1. However, in this case, after encoding *c(t)* into *p*_l_*(t)* such that the phase difference between *p*_l_(*t*) and *p*_2_(*t*) yields *c(t)*, we also encode *c(t)* into the low-frequency signal *s(t)* such that *s(t)* is not random like in scenario 1. We employ the technique used in scenario 2 by utilizing the upper triangular matrix from the Cholesky decomposed covariance matrix to generate *s*_l_(*t*) and *s*_2_(*t*) such that their amplitudes exhibit a time-resolved connectivity of *c(t).* Here, we select a midfrequency band range for *c(t)* in the range of [0.004, 0.008]*Hz* with a mid-coupling amplitude of 0.6. Subsequently, similar to scenarios 1 and 2, *x(t)* and *y(t)* are constructed by summing their respective *s(t)* and *p(t)* signals. As in the previous scenarios, Gaussian noise is added to achieve an SNR of 50dB. We then evaluate Sliding Window Pearson Correlation (SWPC) and phase synchrony between 100-sample pairs of *x(t)* and *y(t)* across multiple window sizes and bandwidths, respectively.

All filtering is done with a Butterworth bandpass filter. The filter’s design aims for a maximum attenuation of 3 decibels in the passband and a minimum of 30 decibels in the stopband. The window sizes used for evaluating SWPC are 15s, 45s, 75s, 105s, and 135s, while the bandwidths used for evaluating phase synchronization are 0.01-0.04Hz, 0.03-0.07Hz, 0.05-0.09Hz, 0.07-0.11Hz, and 0.09-0.13Hz.

### 4.4. Matching parameters

Comparing amplitude-based SWPC with phase-based PS poses challenges due to their inherent differences. SWPC requires a window size, resulting in a shorter time-resolved connectivity time series compared to the original length. PS, employing the Hilbert transform, offers instantaneous measurement, complicating alignment for comparison. Without imposing a window for PS, studying time-resolved similarity becomes impractical. Yet, enforcing a window alters the measure, potentially skewing comparison accuracy. For a low-frequency cutoff of 0.01 Hz, a window size exceeding 88 seconds is recommended to capture the signal’s lowest frequency content (Faghiri, Iraji et al. 2021). To align SWPC with PS, we select two bandwidths (0.01-0.15 Hz and 0.03-0.07 Hz) and corresponding window sizes. Utilizing equation (Error! Bookmark not defined.2) for a low-frequency cutoff of 0.03 Hz, we derive a 30-second window size suited to capture activity within the narrower bandwidth of 0.03-0.07 Hz.

Comparing PS, typically focused on a narrow bandwidth (around 0.03-0.07 Hz) for fMRI data, with SWPC, which operates across all frequencies, presents challenges. While PS limits analysis to a specific frequency range, potentially excluding relevant neuronal information in broader frequency bands, SWPC captures connectivity across all frequencies without such constraints. This discrepancy complicates comparison. Should SWPC be adjusted to match PS’s narrow bandwidth, sacrificing its ability to capture broad connectivity? Alternatively, maintaining SWPC’s full bandwidth may introduce biases. For PS, we adhere to the standard 0.03-0.07 Hz bandwidth. For SWPC, we compute connectivity across the broader BOLD-relevant frequency range (0.01-0.15 Hz) and within the same narrow band used for PS, enabling direct comparison.

Using cosine in PS raises concerns about potential information loss, not always justified in literature. While it offers interpretability by indicating strong synchrony for small phase differences, it masks true differences. For example, cosine of π/4 and 7π/4 both yield approximately 0.7. Despite aligning with SWPC, cosine diminishes information from phase differences.

To align PS with windowed SWPC while preserving PS’s temporal resolution, we use a moving median instead of a moving average. Unlike averages, which blur signal changes, the median retains sharp transitions and timing. We also avoid window-based phase coherence methods for this reason, as they reduce temporal fidelity (Honari, Choe et al. 2020). This approach enables integration of PS and SWPC without compromising PS’s high temporal precision.

### 4.5. Time-resolved comparison between SWPC and PS

For each subject, trFNC is computed using both PS and SWPC methods. These computations are carried out with the specified parameters and the moving median matching technique described above. The computation is conducted across all 53 unique intrinsic brain network pairs obtained from the Neuromark pipeline. To compare the two methods, the brain network pairs are vectorized for each subject, resulting in a *time(window) X reatures* matrix. Here, features represent the vectorized brain network pairs. For each time window of trFNC, the Spearman correlation between SWPC and PS features is computed, yielding time-resolved correlation coefficients between SWPC and PS across features. Spearman correlation was used to compare SWPC and PS because their distributions do not follow a normal distribution (see supplementary material 3) (Hauke and Kossowski 2011). These temporal correlation coefficients depict how SWPC and PS correlate over time for each subject. After obtaining the time-resolved correlation between PS and SWPC for all subjects, these correlation coefficients are separated into five bins based on their values: 0-0.2, 0.2-0.4, 0.4-0.6, 0.6-0.8, and 0.8-1 (no negative values were estimated in our data). These bins illustrate the time windows of trFNC where SWPC and PS are highly correlated and where they are not, according to their discretized correlation bins. For each correlation coefficient bin, the trFNC estimates of SWPC and PS are segregated into their respective bins of correlations, resulting in five sets of trFNC estimates for each method, representing each correlation bin.

To gain insight into the differences between the two methods from the fMRI data, we extract fMRI time points within the windowed trFNC that fall within each Spearman correlation bin. Subsequently, for each timepoint corresponding to a respective correlation bin, we define the same window size applied in trFNC estimations to the fMRI time series centered at that time point. We then computed the power spectral density (PSD) for these windows. Specifically, we conducted a short-time Fourier transform of the fMRI data utilizing the identical window size employed in the trFNC methods. Next, we averaged all power spectra across all time windows, brain networks, and subjects that fall into a given correlation bin. This process resulted in a single PSD representing each correlation bin of interest. This approach allows us to observe how the PSD patterns change based on the correlation between the two methods, potentially providing insight into why these methods may be similar or dissimilar from the data’s power spectrum.

Given the significance of window size selection in short-time power spectrum analysis and its impact on frequency resolution in PSD estimations, we delineate three distinct cases for comparative analysis. In case 1, we utilize the fMRI BOLD relevant frequency range of 0.01-0.15 Hz with a corresponding window size of 88 seconds for SWPC, while a frequency band of 0.03-0.07 Hz is employed for PS, as discussed previously. For case 2, we align the frequency content of both methods by utilizing the 0.03-0.07 Hz frequency band, with a corresponding window size of 30 seconds. To address concerns regarding frequency resolution in case 2, we introduce the third case. In case 3, we maintain the same frequency band as in case 2 but opt for a window size of 88 seconds as in case 1. We conduct all subsequent analyses using these three cases to offer a more comprehensive understanding of their complementary nature, recognizing that no single matching or compensation technique may be adequate to represent the complementary nature of the two methods.

To assess time resolved differences between SWPC and PS across our three scenario cases, we applied the cluster based permutation test to the difference time series (PS–SWPC) for each case. For each case and brain region pair, we computed a paired samples t statistic across subjects on the difference time series, thresholded the resulting t map at uncorrected p < 0.05 to form contiguous supra threshold clusters, and defined each cluster’s mass as the sum of its |t| values. We then generated 10,000 null maxima by randomly sign flipping each subject’s entire difference series, recomputing t maps and cluster masses, and calculating a family wise p value as the proportion of null maxima ≥ the observed cluster mass. Clusters exceeding the 95th percentile of this null distribution (FWER = 0.05) were deemed significant, and the sign of their summed t values indicated whether SWPC or PS dominated over time. These p-values are then corrected using FDR correction across all brain region pairs.

### 4.6. Group analysis with SWPC and PS

To explore the complementary advantages of SWPC and PS, our goal is to leverage the complementary aspects of both methods to gain insights into brain disorders, specifically schizophrenia. Firstly, we compute the average trFNC within each correlation, which is estimated by both SWPC and PS independently. By computing the average connectivity across correlation bins, we can observe the differences in connectivity estimation between the two methods, thereby gaining insights into their average estimations within each correlation bin. For this analysis, we use the fBIRN dataset, focusing specifically on case 1 as described above. In this case, the bandwidth of fMRI for SWPC is set to 0.01 – 0.15 Hz with a window size of 88 seconds, while the bandwidth for PS is set to 0.03 – 0.07 Hz, using a median window of 88s to match it to SWPC.

Furthermore, we extend our analysis of trFNC by incorporating additional steps involving clustering. The aim is to extract recurring patterns (k-means cluster centroids), commonly referred to as brain functional states (Allen, Damaraju et al. 2014). Therefore, after computing the SWPC and PS for the fBIRN dataset, we independently apply k-means clustering to the time-resolved connectivity estimations obtained from each method to identify these recurring patterns.

For this analysis, we continue to use the parameters from case 1: the bandwidth of fMRI for SWPC is set to 0.01 – 0.15 Hz with a window size of 88 seconds, while for PS, the bandwidth is set to 0.03 – 0.07 Hz. However, unlike previous steps, we do not crop the PS estimation using a moving median window. This approach ensures that the results from PS are not compromised by forcing them to match the windowing parameters of SWPC, thereby maintaining the integrity of the PS method and allowing for a fair comparison of the recurring patterns identified by each method.

The k-means clustering is employed with the city-block distance metric, which has been shown to be preferable for high-dimensional data (Aggarwal, Hinneburg and Keim 2001). We chose 4 clusters for PS and SWPC based on the ratio of the within-cluster sum of squared distances (WSS) to the between-cluster sum of squared distances (BSS) (see supplementary material 4 for elbow criteria plots).

The k-means clustering is performed on the full fBIRN dataset, including controls (CN) and schizophrenia (SZ) groups. After clustering, we segregate the clusters based on group membership (CN and SZ) to allow for a direct comparison between the two groups. Two-sample t-tests are further performed on the clusters obtained from PS and SWPC to compare the CN and SZ groups. To control for false positives induced by multiple comparisons, we employ false discovery rate (FDR) correction using the Benjamini-Hochberg procedure (Benjamini and Hochberg 1995). Following common conventions, we employed a significance threshold set at an FDR-adjusted p-value of less than 0.05. This is used for all statistical tests employed in this study.

Furthermore, we computed the average duration that each subject remains within a cluster/state after entering it, known as the mean-dwell time (MDT) (Iraji, Faghiri et al. 2020), and the average percentage of time a subject stays within a cluster/state, known as the fraction rate (FR) (Iraji, Faghiri et al. 2020). To assess the differences between the CN and SZ groups captured by SWPC and PS, we performed a two-sample t-test on the MDT and FR metrics between schizophrenic patients and controls.

Finally, to highlight the complementary nature of SWPC and PS, we analyze how the two methods vary in similarity by quantifying the occupancy of each state obtained from k-means clustering based on the correlation between SWPC and PS. Using the same correlation bins employed in the previous analysis, for each state, we quantify the percentage of each correlation bin that is comprised in that state. In this analysis, a few time points will not belong to any correlation bin in the states obtained from PS since we do not crop PS using a moving median to match it to SWPC before performing k-means clustering. We observed that applying a moving median cropping technique to match PS to SWPC results in a loss of significant information, particularly in the significant differences identified by the PS method in all three cases used in previous analyses (see supplementary material 5). Therefore, to preserve the integrity of PS, we retain these time points. This approach ensures that the PS method’s unique insights are not compromised, allowing for a robust comparison of the complementary aspects of SWPC and PS.

## 5. RESULTS

### 5.1. SWPC and PS using synthesized data

The evaluation of SWPC and PS across various window sizes and frequency bandwidths aims to demonstrate their complementary nature by highlighting their strengths, weaknesses, and similarities. In scenario 1, the ground truth time-resolved connectivity has low magnitude (indicating weak coupling) and is characterized by a high frequency. Figure 3, under scenario 1 reveals consistently low correlation values below 0.4 across all window sizes for the correlations of each 100 sample signal pairs of the SWPC estimations and their respective ground truths, indicating poor and unreliable estimation. Notably, the window size of 15s shows a slightly higher correlation coefficient range between 0.18 to 0.36 compared to other window sizes, which fall below 0.2. This slight improvement in correlation coefficient for the smallest window size is logical, as it allows for more rapid changes in connectivity due to its smaller temporal span. Nonetheless, the estimation remains unreliable. PS demonstrates effective estimation of the ground truth time-resolved connectivity, particularly within the frequency bandwidths of 0.05-0.09Hz, 0.07-0.11Hz, and 0.09-0.13Hz, with correlation coefficients exceeding 0.6, approximately 0.75, and approximately 0.7, respectively. Lower frequency bands for PS fail to effectively estimate the ground truth connectivity, as depicted by the low correlation coefficient distribution under scenario 1 of Figure 3. This outcome is expected since the ground truth connectivity is high frequency. It is intuitive to anticipate higher frequency connectivity within the higher frequencies of the signals rather than within lower frequencies.

**Figure 3.**
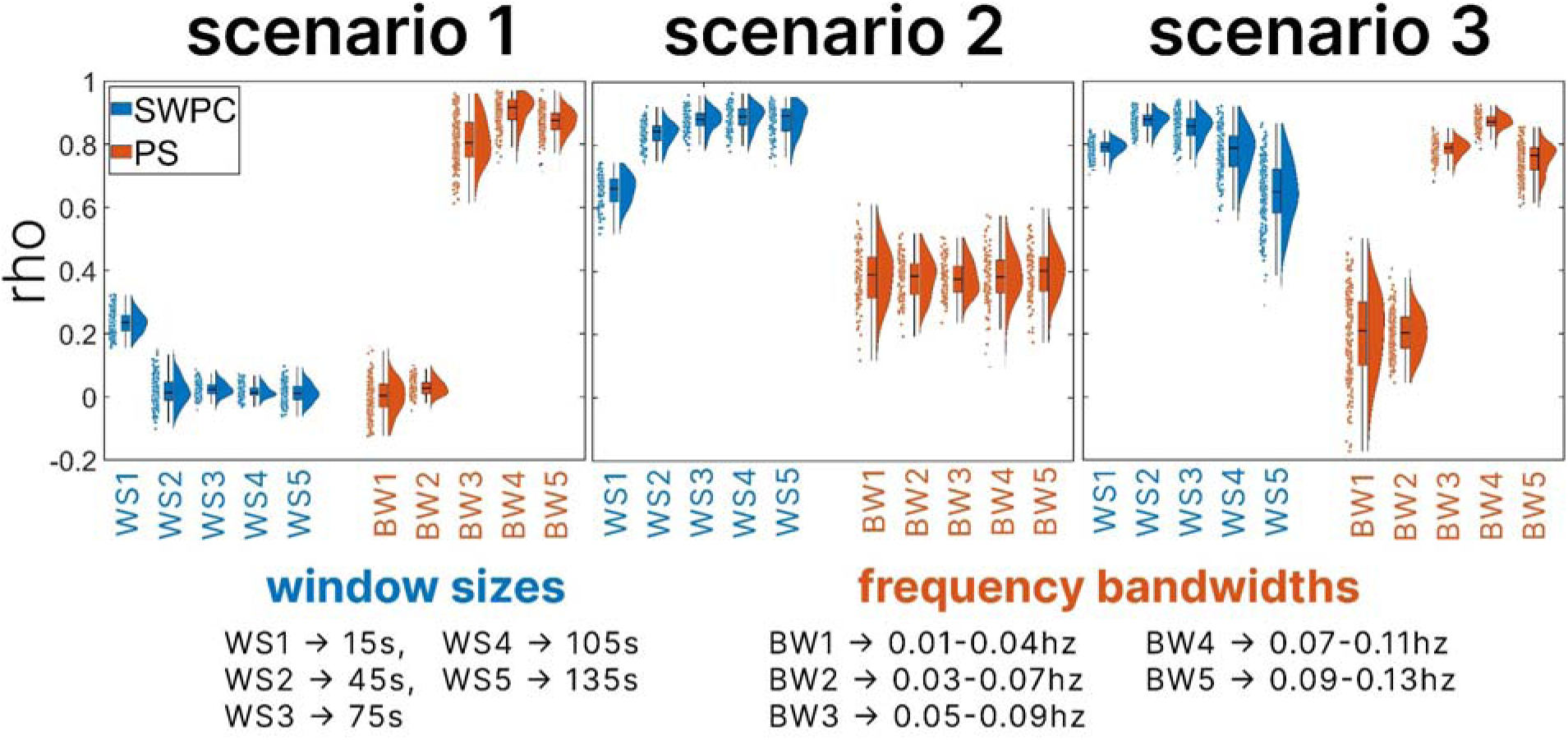
illustrates the results from the simulation analysis of all three scenarios. For each scenario, the correlation between each sample signal pair’s time-resolved connectivity estimations and their respective ground truth time-resolved connectivity is computed. The window sizes used for evaluating SWPC are 15s, 45s, 75s, 105s, and 135s, while the bandwidths used for evaluating phase synchronization are 0.01-0.04Hz, 0.03-0.07Hz, 0.05-0.09Hz, 0.07-0.11Hz, and 0.09-0.13Hz. The boxplots, kernel density, and scatter plots of correlation coefficients across all samples for each scenario are displayed, with SWPC results shown in blue and PS results shown in orange. Note that PS performs better in scenario 1, and SWPC performs better in scenario 2 emphasizing the complementary nature of the two approaches.

In scenario 2, where the time-resolved connectivity exhibits high magnitude (indicating strong coupling) and low frequency (reflecting stable connectivity), higher correlation coefficients are observed between the SWPC estimations and their respective ground truths across all window sizes in Figure 3. However, the 15s window size shows a slightly lower distribution of correlation coefficients compared to other window sizes. Despite this slight decrease, the correlation coefficients against the PS estimations remain higher for the 15s window size under scenario 2 in Figure 3. SWPC demonstrates a reliable estimation of high magnitude, low-frequency time-resolved connectivity. In contrast, the correlation coefficients between PS estimations and the ground truth are more spread out, centered around 0.4, indicating an unreliable estimation of strong low-frequency time-resolved connectivity.

Scenario 3 was designed to depict a scenario where the time-resolved connectivity is encoded in the amplitude of the signal pairs rather than the phase, with the characteristics of the time-resolved connectivity representing a compromise between scenario 1 and 2. This entails mid-strength and mid-frequency time-resolved connectivity. Both SWPC and PS estimations demonstrate high correlation coefficients when compared with their respective ground truths, particularly for window sizes of 15s, 45s, and 75s, as well as frequency bandwidths of 0.05-0.09Hz, 0.07-0.11Hz, and 0.09-0.13Hz, respectively.

The simulation results compared using the root mean squared error between the SWPC and PS estimations with their respective ground truths is given in supplementary material 6.

### 5.2. Time-resolved comparison of SWPC and PS

The comparative analysis between SWPC and PS methodologies, looking into the distinctions between these approaches and their relationship to the dynamic activity of brain networks, is displayed in Figure 4. Figure 4 illustrates the outcomes of temporal correlation assessments between SWPC and PS across pairs of brain networks and subjects. The analysis is performed for three distinct cases: Case one employs a bandwidth of 0.01 to 0.15 Hz for SWPC with a window size of 88 seconds, while cases two and three utilize a bandwidth of 0.03 to 0.07 Hz for SWPC. However, case two employs a window size of 30 seconds, corresponding to the low-frequency cutoff of the fMRI data, while case three retains the window size of 88 seconds. For all cases, PS utilizes the fMRI bandwidth of 0.03 to 0.07 Hz. The PDFs of the time-resolved correlation coefficients between SWPC and PS are depicted in Figure 4 (A) for all cases and both the fBIRN and HCP dataset. Notably, the PDFs derived from both the fBIRN and HCP datasets exhibit very high similarity, with substantial overlap within each case.

**Figure 4.**
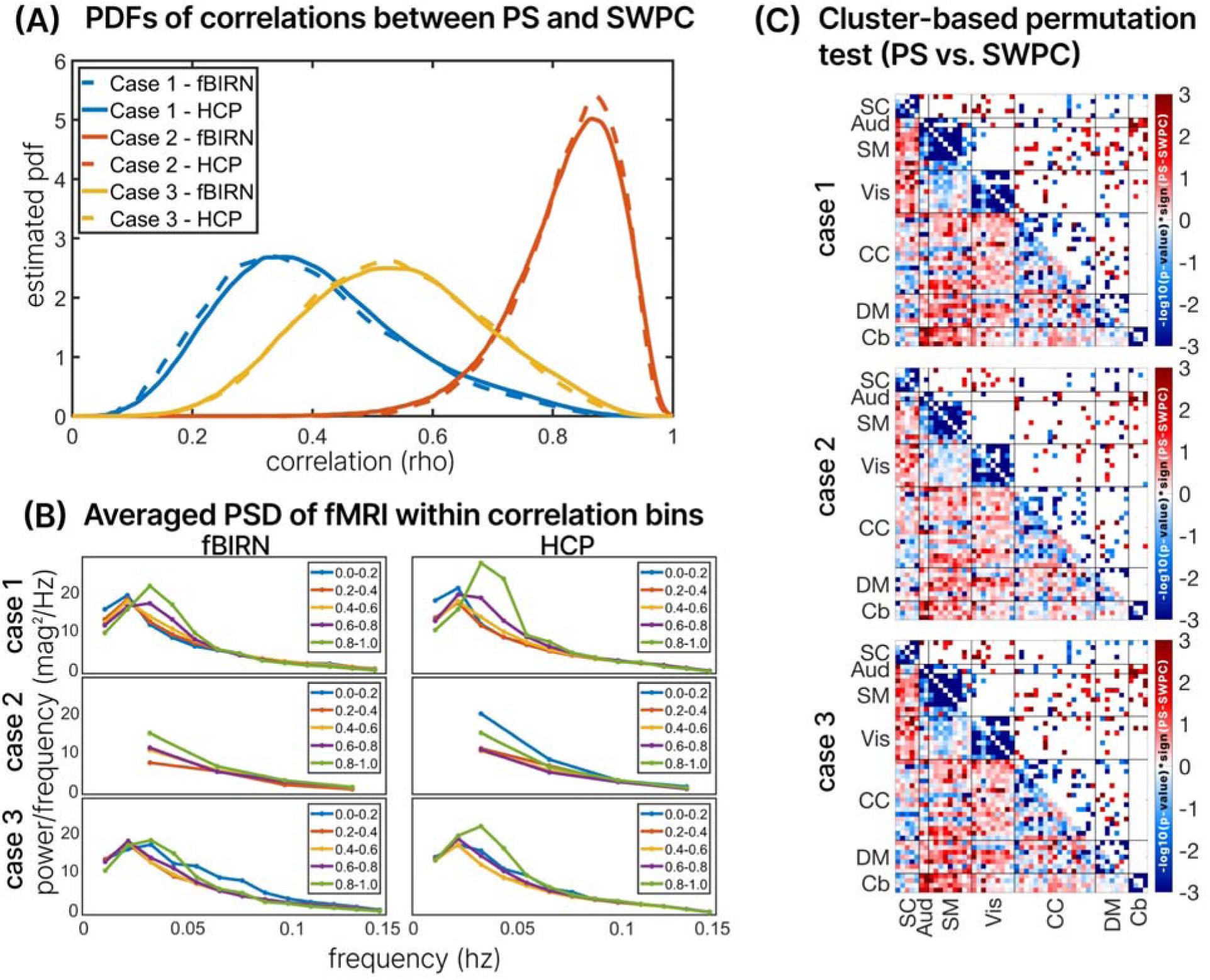
presents results from segregating Phase Synchrony (PS) and Sliding Window Pearson Correlation (SWPC) into five temporal correlation bins (0-0.2, 0.2-0.4, 0.4-0.6, 0.6-0.8, and 0.8-1) across three cases. In Case 1, SWPC has a bandwidth of 0.01-0.15 Hz with a window size of 88 seconds, while PS uses 0.03-0.07 Hz. In Case 2, both SWPC and PS use a bandwidth of 0.03-0.07 Hz with a window size of 30 seconds. In Case 3, both methods utilize a bandwidth of 0.03-0.07 Hz, but the window size is 88 seconds. (A) depicts the probability density function (PDF) of SWPC-PS correlation coefficients for each case. Additionally, the Power Spectral Density (PSD) of the fMRI data from HCP and fBIRN datasets across varying correlation bins is shown in (B). The averaged PSD across windows of the fMRI time series, brain networks, and subjects corresponding to each correlation bin is displayed for each case. This highlights that SWPC and PS are most similar (within high correlation bins) when the frequency range accessed by both approaches is close, particularly when the fMRI data exhibits a strong dominant frequency. (C) displays the results from a cluster-based permutation test using 10,000 samples to build a null model to test the difference between SWPC and PS. The lower triangle shows the -log10*(p-values)*sign(t-values) while the upper triangles show the significantly thresholded version (alpha = 0.05).

In the context of case one, the time-resolved correlation between PS and SWPC is notably lower compared to the other cases, which aligns with the different frequency content accessible to each method (0.01 to 0.15 Hz for SWPC and 0.03 to 0.07 Hz for PS). Conversely, in case two, where the frequency contents are the same, the correlation between PS and SWPC is significantly higher, as expected. Finally, in case three, the time-resolved correlation coefficients fall between those of case one and case two, indicating an intermediary relationship between SWPC and PS correlations. These findings reveal the nuanced interplay between the parameters and their impact on the comparative analysis of PS and SWPC in relation to trFNC of the brain.

In Figure 4(B), the averaged PSDs corresponding to the correlation bins are presented for each case. In case 1, the PSD patterns exhibit consistency between the fBIRN and HCP datasets. Notably, for bins indicating a high correlation between SWPC and PS, there is a discernible peak around 0.033 Hz in the fMRI data, particularly within correlation bins of 0.8-1 and 0.6-0.8. Conversely, for bins indicating lower correlations between SWPC and PS, a peak in PSD is observed at 0.022 Hz of the fMRI data. These patterns persist across datasets. In case 2, the PSD patterns show slight discrepancies between the fBIRN and HCP datasets. Specifically, in the HCP dataset, there exists a PSD plot for the lowest correlation bin, which is absent in the fBIRN dataset. Upon examining the PDFs in Figure 4(A) for case 2, it becomes evident that very few subjects fall within the low correlation bins. Consequently, there are minimal subjects within correlation bins below 0.6, rendering the PSD estimations below 0.6 unreliable. The observed high power at a frequency around 0.033 Hz in the correlation bin of 0-0.2 for case two in the HCP dataset may be attributed to a small subject size, as there are no subjects in the fBIRN dataset for this correlation bin. This poses a challenge in interpreting the results of case 2, especially considering the window size of 30 seconds, which results in poorer frequency resolution. However, for the reliable bins in case 2, namely 0.8-1 and 0.6-0.8, the patterns remain consistent across both datasets: higher correlation bins correspond to higher peak power at 0.033 Hz.

In case 3, where the window size is 88 seconds and the frequency content matches in SWPC and PS (0.03-0.07hz), the PSD patterns exhibit consistency between the fBIRN and HCP datasets. Notably, for the highest correlation bin (0.8-1.0), there is a peak of PSD at 0.033 Hz, mirroring the findings observed in case 1. When examining lower correlation bins—specifically, bins of 0.6-0.8, 0.4-0.6, and 0.2-0.4—a peak at 0.022 Hz in the PS is evident, with the highest correlation bin among these three (0.6-0.8) exhibiting a slightly higher peak. This consistency is observed across both datasets. However, multiple peaks at various frequencies are observed for the lowest correlation bin, 0-0.2. While this pattern is consistent with both datasets, it is more prominent in the fBIRN dataset.

To further demonstrate the complementary nature of SWPC and PS in fMRI connectivity, we applied a cluster-based permutation test under the three cases mentioned above. In every case, the method specific differences exceeded the null distribution: SWPC consistently produced stronger within domain connectivity across contiguous time segments, whereas PS showed significantly elevated between domain connectivity.

### 5.3. Group differences with SWPC and PS

The averaged trFNC across subjects and time (windows) for each correlation bin is computed on the fBIRN dataset to observe how the functional connectivity estimated using SWPC and PS varies as a function of the similarity between the two methods. The results are depicted in Figure 5. Specifically, Figure 5(A) shows the averaged trFNC within each correlation bin, segmented into 5 segments from correlation coefficients from 0 to 1. The trFNC patterns exhibit strongly connected brain networks within the bins where correlation is highest, such as in the 0.8-1 and 0.6-0.8 correlation coefficient bins. For the lower correlation bins, both SWPC and PS showcase reduced levels of connectivity estimations, with PS exhibiting lower levels of connectivity patterns compared to SWPC. This indicates that the dissimilarities between SWPC and PS in estimating trFNC are more pronounced in scenarios with low connectivity patterns. In this analysis, the frequency band for SWPC is 0.01-0.15 Hz with a window size of 88 seconds, while the frequency band for PS is 0.03-0.07 Hz with a cropping moving median to match PS to SWPC.

**Figure 5.**
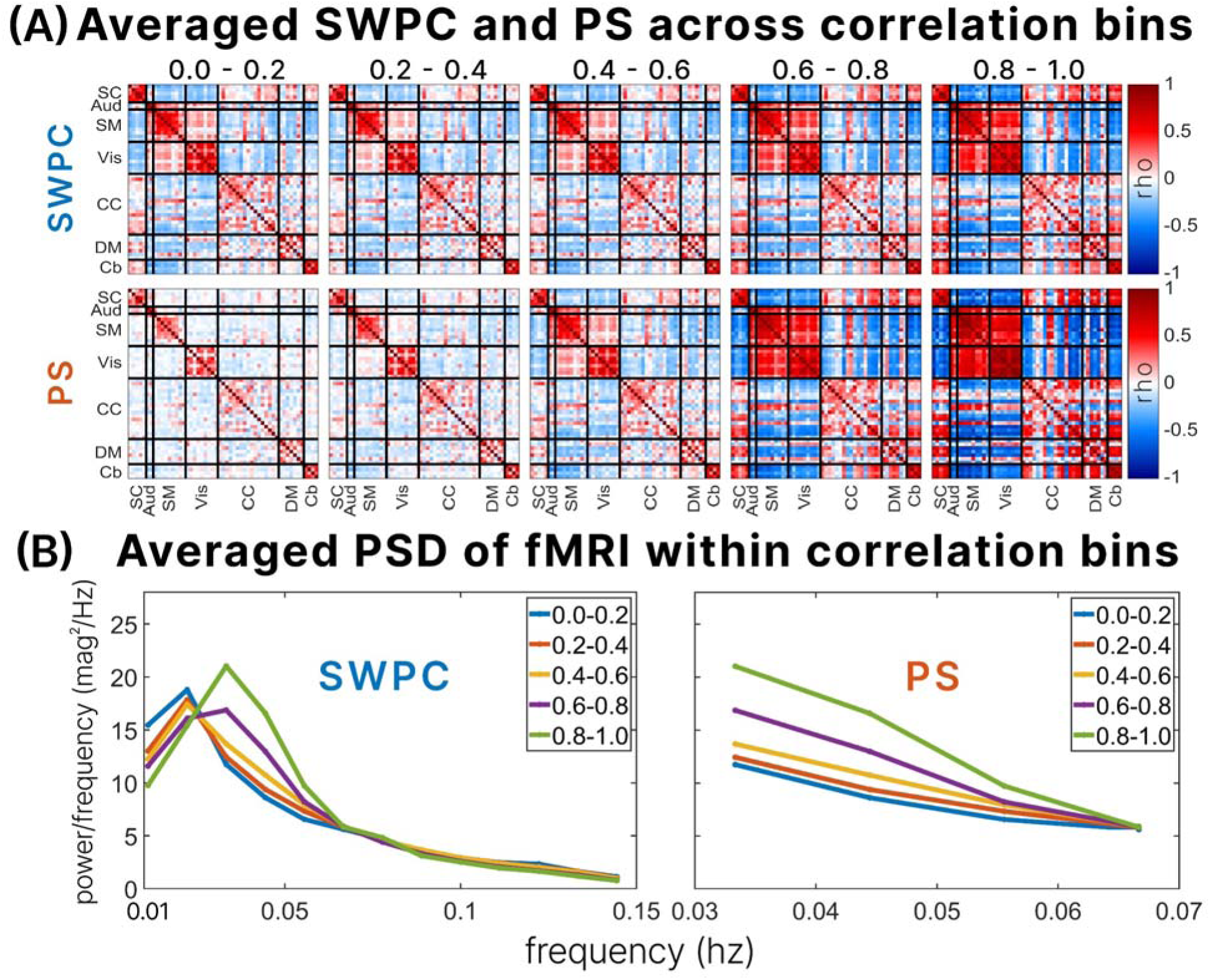
shows the averaged trFNC across brain network pairs within each correlation bin, depicting the similarity/dissimilarity between the trFNC estimated by SWPC and PS based on Spearman correlation. For each correlation bin, the average trFNC is displayed in (A), while the average power spectral density (PSD) across all brain networks and subjects is shown in panel (B). For each time point within a correlation bin, a window is defined centered at that time point of the fMRI data across all brain networks with a window size of 88 seconds. The PSD is then computed for these windows. All PSDs are averaged across all subjects, time/windows, and brain networks within each correlation bin to obtain a single PSD corresponding to each correlation bin. Considering that PS utilizes a bandwidth of 0.03 - 0.07 Hz, we display the PSD within this range in panel (B) for PS. Note that the lower the correlation between SWPC and PS, the less connected the trFNC estimations derived from SWPC and PS are, with this effect being more pronounced for PS. Also, when the correlation between SWPC and PS is strong, the dominant frequency of the fMRI data accessed by SWPC closely matches that of PS, which is 0.033 Hz. For lower correlations, the dominant frequency of the fMRI data accessed by SWPC is around 0.022 Hz, indicating that the connectivity varies across different frequency bands as accessed by the two methods.

The PSD of the fBIRN fMRI frequency band accessed by SWPC and PS within each correlation bin is displayed in Figure 5 (B). The dominant frequency component of the fBIRN fMRI data across subjects and brain networks is 0.033 Hz for all correlation bins in PS, but it shows reduced power as the correlation bins decrease. The high power in high correlation bins at the dominant frequency indicates high synchronization at the 0.033 Hz frequency, suggesting a form of oscillatory behavior, which explains the strongly connected patterns observed within the high correlation bins in Figure 5 (A) for PS.

For SWPC, the dominant frequency within the highest correlation bin is around 0.033 Hz, which implies matched information between the two methods considering this is the same dominant frequency observed in the case of using PS. However, for lower correlation bins, the averaged PSD accessed by SWPC still shows oscillatory properties, but with a dominant frequency at 0.22 Hz instead. Considering that PS does not have access to this frequency range, it is expected that at a dominant frequency of 0.22 Hz, the similarities between SWPC and PS will be reduced.

These observations suggest that the differences between SWPC and PS are more evident in lower connectivity states and are also due to the different frequency bands utilized by the two methods. This highlights the complementary nature of SWPC and PS in capturing different aspects of brain connectivity, with each method providing unique insights that the other might miss.

To further explore the complementary nature of SWPC and PS, we perform a group analysis on the states estimated by the two methods between SZ and CN groups using the fBIRN dataset. Figure 6 displays the results from the group analysis, showcasing the complementary functional relevance to brain disorder between the two methods. Figure 6 (A) shows the cluster centroids of SZ (upper triangle) and CN (lower triangle) for SWPC and PS.

**Figure 6.**
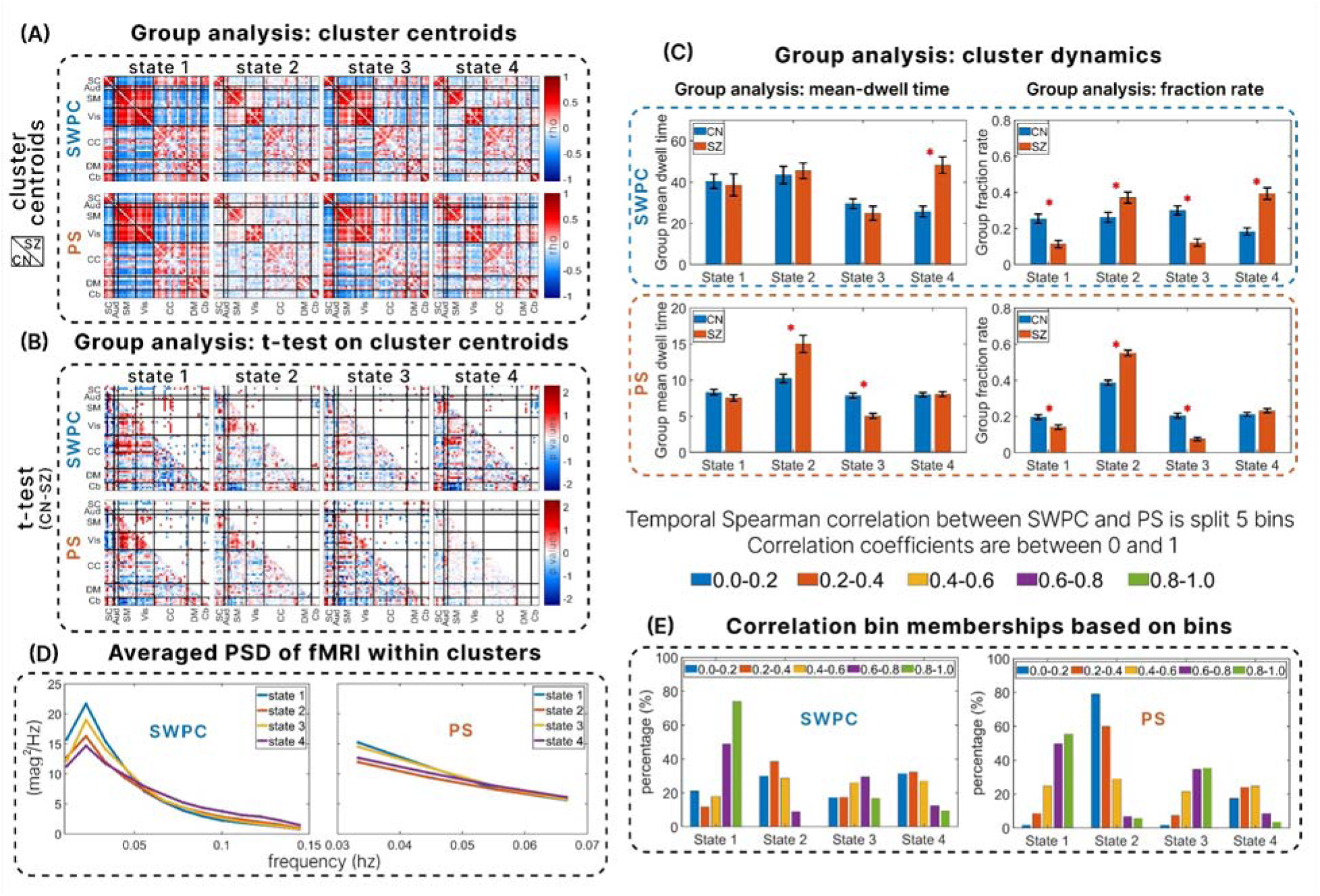
illustrates the group analysis results obtained from computing k-means clustering on SWPC and PS. The states derived from k-means clustering for SWPC and PS are depicted in (A). In these matrices, the upper triangle represents the states of brain network pairs for the schizophrenic (SZ) group, while the lower triangle represents the states of brain network pairs for the control (CN) group. (B) shows the results of a two-sample t-test performed on the estimated states to identify significant differences between SZ and CN groups in the brain network pairs within the states for SWPC and PS independently. The lower triangle displays the -log(pvalue) * sign(CN - SZ), indicating both the direction and significance of the differences. The upper triangle shows the thresholded p-value, with a threshold of less than 0.05. Note how the significant differences identified by SWPC and PS vary across states, highlighting their complementary nature. (C) displays the mean-dwell time (MDT) and fraction rate (FR) of SWPC and PS between SZ and CN groups, with significant differences indicated by an asterisk based on a p-value threshold of less than 0.05. (D) depicts the average power spectral density (PSD) of the fMRI data utilized by SWPC (0.01 - 0.15hz) and PS (0.03-0.07hz), averaged across subjects and time within each cluster/state. Note that MDT of SWPC finds significant differences in states 2 and 3, while PS identifies significant differences in state 4, indicating the complementary nature of the information they provide. (E) shows the percentage of membership of each correlation bin within a cluster/state for both SWPC and PS, indicating the percentage each correlation bin forms a cluster.

In the states derived from k-means clustering of SWPC and PS, there are two high connectivity/anti-connectivity states and two low connectivity/anti-connectivity states for both methods, as shown in Figure 6(A). These states are supported by the fMRI power spectral density (PSD) accessed by each method, illustrated in Figure 6(D). Low power of the dominant frequencies is observed in states 2 and 4, emphasizing these as the states with the least level of synchrony across brain network pairs. Given that the difference between the two methods is more evident in low connectivity states, we expect these low connectivity states for both SWPC and PS to be occupied by a larger percentage of low correlation bins, specifically the 0-0.2 and 0.2-0.4 correlation coefficient bins. Figure 6 (E) confirms this expectation, showing that states 2 and 4 have the highest percentage occupancy of those correlation bins for both PS and SWPC, further emphasizing these as the least connected states. Conversely, states 1 and 3 exhibit the highly connected brain network paired states. This is also supported by the PSDs of PS and SWPC, which exhibit the highest power at the dominant frequencies in Figure 6(D). Additionally, in Figure 6(E), states 1 and 3 show the highest percentage occupancy of the highest correlation bins, specifically in the 0.6-0.8 and 0.8-1 bins.

Figure 6(B) presents the results of a two-sample t-test with FDR correction. The lower triangle indicates -*log(pvalue) * sign(t - score),* providing both the direction and a continuous representation of the significant differences. The upper triangle shows the thresholded FDR-corrected p-values, also indicating the direction of the differences as in the lower triangle.

These significant differences between SZ and CN, as captured by PS and SWPC, are complementary. For example, in the most connected state, state 1 in Figure 6(B), PS captures significant differences between the sensory-motor and visual network pairs, which are missed by SWPC. This indicates that controls exhibit significantly higher synchronization than schizophrenic groups between sensory-motor and visual networks. Conversely, in state 1 of SWPC, there is a significant difference between SZ and CN in the connectivity between sensory-motor networks and cognitive networks, which is missed by PS. This indicates that CNs exhibit significantly higher correlations than SZ groups between their sensory-motor and cognitive networks.

The mean dwell time and fraction rate between SZ and CN further emphasize the complementary nature of these methods, as depicted in Figure 6(C). The asterisks in Figure 6(C) indicate significant p-values between the MDT and FR of SZ and CN using the two methods, with a threshold of less than 0.05. PS captures significant differences between SZ and CN groups in MDT for states 2 and 3. As shown in Figure 6(A), state 2 is a less synchronized state, while state 3 is a high synchronization/anti-synchronization state. These significant differences are not detected by SWPC in its corresponding states. Conversely, SWPC captures significant differences between SZ and CN groups in MDT for state 4, another less connected state, which PS does not detect. Schizophrenia has been widely associated with less connected brain networks (Lynall, Bassett et al. 2010, Nejad, Ebdrup et al. 2012, Damaraju, Allen et al. 2014, Rabany, Brocke et al. 2019). Consequently, we expect significant differences between SZ and CN groups in the MDT of the less synchronized/connected states. According to Figure 6(A), state 4 is the least connected state in SWPC and shows the lowest dominant frequency in Figure 6(D). Thus, it is anticipated that schizophrenic patients are more likely to remain in state 4 once they enter it, compared to controls in SWPC.

Interestingly, in PS, the least synchronized state according to Figure 6(A) and the one with the lowest dominant frequency in Figure 6(D), is state 2. As expected, schizophrenic patients are more likely to stay in this state longer than controls. The differences in the least connected states identified by SWPC and PS are noteworthy. Although both methods identify a disconnected state, these states are different, suggesting that schizophrenic patients exhibit different types of functionally disconnected brain network patterns detected by PS and SWPC, particularly when examining the mean dwell time of trFNC.

In terms of FR, SWPC captures significant differences between SZ and CN in all four states, while PS captures significant differences in the first three states. In state 4, SWPC exhibits the least connected brain networks, whereas the PS counterpart shows some modular positive, relatively low synchrony patterns between sub-cortical, auditory, and sensory-motor brain networks. State 4 of PS is primarily occupied by correlation bins 0.2-0.4 and 0.4-0.6, as shown in Figure 6(E). This implies that, for the mid-range similarity between SWPC and PS, specifically the correlation bins 0.2-0.4 and 0.4-0.6, PS identifies some unique patterns that SWPC does not detect. However, while these patterns are interesting, they do not provide any significant difference between schizophrenic patients and controls in terms of FR.

These results further highlight the complementary insights provided by each method. Each method reveals unique aspects of connectivity dynamics, with PS being more sensitive to certain states and SWPC to others, thereby providing a more comprehensive understanding when used together.

## 6. DISCUSSION

From a communication theory perspective, phase synchronization can be viewed as a phase demodulation technique (Wang and Da 2012), while sliding window Pearson correlation can be seen as an amplitude demodulation technique (Faghiri, Iraji et al. 2022) as discussed extensively in the background section. These two techniques are fundamentally different and offer complementary strengths and limitations. This distinction emphasizes that PS and SWPC, as time-resolved functional network connectivity measures, capture different information: PS extracts shared information through the phase of the BOLD activity, whereas SWPC captures shared information through the amplitude of the BOLD activity.

Moreover, these methods differ in their ability to capture various kinds of connectivity between a pair of signals. Our simulations demonstrate that PS more accurately estimates low-amplitude fast-varying connectivity between narrow band signal pairs, as shown in scenario 1 of Figure 3 and consistent with previous findings (Fagerholm, Moran et al. 2018, Lee and Sepulchre 2024). On the other hand, high-amplitude slow-varying connectivity between broadband activity signal pairs is better estimated by SWPC, as illustrated in scenario 2 and is consistent with previous findings (Scott, Fagerholm et al. 2014, Fagerholm, Scott et al. 2016, Fagerholm, Moran et al. 2020). These findings highlight the differences in the types of connectivity each method captures.

While the distinction between the two methods is clear, it is not uncommon to expect phase and amplitude information to be similar at certain times and in specific scenarios. This leads to an interest in comparing SWPC and PS in the context of trFNC. However, comparing these two measures is not straightforward due to their differing nature. For instance, PS operates as an instantaneous measure, while SWPC functions as a window-based measure. In our study, we compare these two methods across several cases, across window sizes and bandwidths in fMRI data, acknowledging that no single comparison case is the best. We aim to link differences between the two methods with the power spectral density estimation of fMRI data. This approach allows us to understand variations in trFNC by analyzing the spectral properties of fMRI data across Spearman correlation bins corresponding to the two methods.

In case 1, we use standard practices of an 88-second window and a frequency band of 0.01-0.15 Hz for SWPC, and match PS using a median window of the same window and a frequency band of 0.03-0.07 Hz. To address the frequency range mismatch between SWPC and PS, we use a frequency band of 0.03-0.07 Hz for SWPC with a corresponding window size of 30 seconds in case 2. However, the 30-second window significantly reduces the frequency resolution of the PSD estimation. To counter this, we create Case 3 from Case 2 by using an 88-second window instead of 30 seconds. While these cases aim to balance the compromises of the two methods, we acknowledge that neither of them is the most appropriate case for comparison. Nonetheless, we find consistency in the PSD estimations across all cases and two different fMRI datasets in relation to the correlation bins between SWPC and PS.

As shown in Figure 4(A), the estimated PDF of the Spearman correlation between SWPC and PS is consistent across the fBIRN and HCP datasets, suggesting that the time-varying correlations between SWPC and PS are not biased by a single dataset. The correlation between SWPC and PS is highest in Case 2, which is expected since a smaller window size for SWPC aligns more closely with the instantaneous nature of PS. However, it is important to note that amplitude correlations are best suited for amplitude estimation (Van Den Heuvel and Pol 2010, Engel, Gerloff et al. 2013, Fu, Tu et al. 2018, Mostame and Sadaghiani 2020), which requires a low-pass filter to extract the relevant slow-varying amplitude of the signal (Faghiri, Iraji et al. 2021), as done in amplitude demodulation (Haykin and Moher 1989, Proakis, Salehi et al. 1994) and consequently by SWPC. Therefore, while SWPC closely matches PS in Case 2, SWPC is not reliable in this case due the conceptual inconsistency and also to the reduced statistical power of correlation estimations (Schönbrodt and Perugini 2013).

The PSD estimations of the fMRI data across correlation bins show consistency with the concept of phase and amplitude being synced under strong oscillatory conditions (Fagerholm, Moran et al. 2020). Intuitively, if two sine waves have the same frequency and phase shifts, phase synchronization and amplitude correlations will be synchronized. If two signals are strongly coupled, their dominant frequencies tend to align (Rosenblum, Pikovsky and Kurths 1996, Boccaletti, Kurths et al. 2002), giving rise to oscillations at the same or nearly the same frequencies (oscillatory conditions) (Lowet, Roberts et al. 2016, Sivarajah, Steinbacher et al. 2019, He, Xu et al. 2022). This oscillatory behavior can be observed when the average power spectral density across all brain networks reflects a dominant frequency. Consequently, we expect SWPC and PS to be closely matched when the fMRI PSD across all subjects and brain networks is closer to an oscillatory condition, which is observed in the higher correlation bins (0.6-0.8 and 0.8-1.0). This is observed in Figure 4(B) across all cases and both datasets, especially for the correlation bin 0.8-1.0, except in Case 2 of the HCP dataset where the lowest correlation bin (0.0-0.2) yields the highest power of the dominant frequency in the PSD estimation, implying strong oscillatory behavior. This anomaly can be explained by the low frequency resolution in that case, making the PSDs unreliable. Another possible reason could be drawn from scenario 2 of our simulation results, which shows that SWPC outperforms PS when connectivity is encoded in the amplitude of the signals (Van Den Heuvel and Pol 2010, Engel, Gerloff et al. 2013, Mostame and Sadaghiani 2020) with a high amplitude of connectivity (Fagerholm, Moran et al. 2020). This suggests that the strong oscillatory component is encoded in the amplitude of the brain networks at those times. However, this is not observed in the fBIRN dataset. In the fBIRN dataset, no samples exist within the 0-0.2 correlation bin in case 2 (Figure 4A) and thus, have no corresponding PSD estimation (Figure 4B). Due to this, the reason for the strong oscillatory component being encoded in the amplitude in the HCP dataset remains conjectural since it cannot be validated in the fBIRN dataset.

To demonstrate that the methods’ differences extend beyond their spectral filters and reflect the fundamental distinctions described in our background, we applied the cluster-based permutation test across all three scenarios. In every case, SWPC showed significantly stronger within-domain connectivity (especially within the sensory networks), whereas PS exhibited significantly greater between-domain connectivity, underscoring their complementary nature, Figure 4 (C). Supporting evidence for this pattern already exists in electrophysiology. Hipp et al. reported that amplitude-envelope correlations are strongest within sensory networks, whereas phase synchrony is comparatively weaker across them (Hipp, Hawellek et al. 2012). Florin and Baillet found that phase-based coupling facilitates long-range, cross-system communication, complementing more local amplitude interactions (Florin and Baillet 2015). Finally, Siems and Siegel showed that amplitude- and phase-coupling form dissociable networks, with amplitude coupling predominantly local and phase coupling more distributed (Siems and Siegel 2020).

Another reason for the difference between SWPC and PS is the mismatch of frequency bands: SWPC utilizes a range of 0.01-0.15 Hz, while PS utilizes 0.03-0.07 Hz. This discrepancy is evident in Figure 5. Figure 5 shows the average trFNC and average PSD of the fBIRN fMRI dataset across correlation bins between SWPC and PS.

From Figure 5(A), the higher the correlation bin between the methods, the more connected or synchronized the brain network pairs are for both methods. Conversely, the lower the correlation bin, the less connected the brain network pairs appear, especially in PS. The low level of connectivity of brain networks in the low correlation bins is supported by the estimated PSD within these bins. In Figure 5(B), it is evident that the lower the correlation bin, the lower the power of the dominant frequency at 0.033 Hz, implying that lower correlation bins correspond to less oscillatory properties of the fMRI data. This aligns with the observation that PS shows a lower level of synchronization in lower correlation bins. For lower correlation bins, the fMRI data accessed by SWPC shifts from the 0.03 Hz frequency to lower dominant frequencies around 0.022 Hz. Despite this shift, the PSDs for these lower correlation bins still exhibit oscillatory properties, explaining why low correlation bins in SWPC still show a relatively higher level of connectedness in Figure 5(A). This underscores one of the reasons SWPC and PS differ: the mismatch of frequency bands. This observation is also consistent with the notion that functional connectivity varies across different frequency bands (Luo, He et al. 2020, Faghiri, Iraji et al. 2021, Ziaeemehr and Valizadeh 2021, Li, Cheng et al. 2022, Kajimura, Margulies and Smallwood 2023, Neşe, Harı et al. 2024).

To explore the clinical relevance of the disparities between SWPC and PS, we conducted k-means clustering on the trFNC estimated by SWPC and PS independently to obtain functional connectivity states as displayed in Figure 6(A). The composition of each correlation bin in each cluster/state is displayed in Figure 6(E). One notable distinction between SWPC and PS is evident in state 4. Despite being a low-connected state, PS exhibits an observable modular structure more prominently than SWPC. PS detects low levels of synchronization between sub-cortical, auditory, and sensory-motor brain networks, which are not detected by SWPC. Connecting this observation to the similarities between the two methods, it can be seen from Figure 6(E) that state 4 has a high percentage of mid-range correlation bins for PS, specifically the 0.2-0.4 and 0.4-0.6 bins. This highlights that at mid-range correlations between the two methods, PS identifies synchrony patterns that are missed by SWPC. Considering that this state is a low-connected/synchronized state, a plausible reason why this pattern is found for PS could be drawn from our simulation analysis, which suggests that PS is more appropriate for capturing low-amplitude time-resolved connectivity than SWPC (Fagerholm, Moran et al. 2018, Lee and Sepulchre 2024), as demonstrated in scenario 1 of Figure 3.

The state with the least connected brain networks for SWPC is state 4, while for PS it is state 2, as shown in Figure 6(A) and validated by the PSD estimation of the fBIRN fMRI dataset across the states in Figure 6(D). The state with the lowest PSD at the dominant frequency is shown as state 4 (purple plot) for SWPC and as state 2 (red plot) for PS. One striking observation is that the least connected states manifest differently in SWPC and PS, highlighting their complementarity. The patterns of state 4 in SWPC are characterized by low anti-correlations between visual networks and subcortical, auditory, and sensory-motor networks. In contrast, state 2 in PS is characterized by low synchronization between the same brain networks. We detect significant differences between schizophrenia and control groups in state 2 of PS and state 4 of SWPC, with SZ groups showing a higher mean-dwell time in these states than CN groups, as shown in Figure 6 (C). This finding aligns with previous studies indicating that individuals with schizophrenia are more associated with low connectivity levels (Friston and Frith 1995, Friston 1998, Kim, Burge et al. 2008, Lynall, Bassett et al. 2010, Pettersson-Yeo, Allen et al. 2011, Nejad, Ebdrup et al. 2012, Williamson and Allman 2012, Damaraju, Allen et al. 2014, Miller, Yaesoubi et al. 2016, Iraji, Deramus et al. 2019, Iraji, Fu et al. 2019). These results suggest that while SZ groups tend to stay longer in the least connected/synchronized states (Damaraju, Allen et al. 2014, Du, Pearlson et al. 2016, Fu, Caprihan et al. 2019, Faghiri, Iraji et al. 2020, Weber, Johnsen et al. 2020, Fu, Iraji et al. 2021, Hyatt, Wexler et al. 2022, Bostami, Lewis et al. 2024), there are multiple low connected/synchronized states associated with schizophrenia. One state is characterized by low anti-correlations between visual networks and subcortical, auditory, and sensory-motor networks, as detected by SWPC (Damaraju, Allen et al. 2014, Du, Pearlson et al. 2016, Faghiri, Yang et al. 2023). The other state is characterized by low positive synchronization between the same brain networks, as detected by PS in Figure 6(A).This dual pattern of low connectivity states highlights the complexity of functional connectivity alterations in schizophrenia and underscores the importance of using both SWPC and PS to capture different aspects of brain network dysfunction in this disorder.

Another clinically relevant difference between SWPC and PS can be observed in state 3. In state 3 of SWPC, most cognitive control networks exhibit anti-correlations with sensory-motor and visual networks, while in PS, there exist positive synchronization among some networks within those domains, as depicted in Figure 6(A). We observe more significant differences between schizophrenia and control groups in state 3 of PS than we detect in state 3 of SWPC, as illustrated in Figure 6(B). For instance, CN exhibits significantly higher synchronization between some sensory-motor networks and visual networks, indicated by the red dots in state 3 of PS in Figure 6(B). Additionally, we detect significant differences between SZ and CN groups in state 3 of PS, with CN groups showing a higher mean-dwell time in this state than SZ groups, as shown in Figure 6(C), an observation undetected by SWPC. According to Figure 6(E), state 3 constitutes declining percentages of correlation bins from high to low. This pattern is similar to state 1 and validated by Figure 6(D), where states 1 and 2 are the most connected states, with state 1 being slightly higher than state 3. This demonstrates that even for highly connected/modular states and states mostly characterized by high correlation between SWPC and PS, the clinically relevant information is complementary. This observation is further made evident in state 1 for both methods, as the significant differences between SZ and CN groups are complementary, as shown in Figure 6(B). SWPC indicates that in state 1, CN groups exhibit stronger correlations between sensory-motor networks and cognitive control networks, undetected by PS, while PS identifies stronger synchronization between sensory-motor networks and visual networks, undetected by SWPC.

Neurobiologically, the observation that SWPC isolates a state dominated by strong anti-correlations among visual, auditory, sensorimotor and sub-cortical circuits while PS uncovers a parallel state with low-amplitude, yet phase-coherent positive couplings may suggest that the two estimators are capturing complementary facets of schizophrenia-related dysconnectivity. SWPC appears particularly sensitive to periods of pathological segregation of sensory systems, which could be rooted in abnormal thalamo-cortical gating and excessive network antagonism(Damaraju, Allen et al. 2014, Anticevic, Haut et al. 2015). By contrast, the subtle hypoconnected but synchronous regime detected only with PS may reflect the excitatory– inhibitory imbalance predicted by N-methyl-D-aspartate receptor hypofunction models, where reduced excitatory drive lowers coupling *amplitude* without disturbing oscillatory *timing* (Anticevic, Gancsos et al. 2012, Jadi and Sejnowski 2014). Recognizing that patients may cycle between these two regimes may help explain why each method reveals distinct dwell-time abnormalities and underscores the value of analyzing SWPC and PS together to capture the full spectrum of functional network disruption.

Based on our results, it is strongly evident that the two methods are complementary, each with their own strengths and weaknesses. The suggestion to use PS instead of SWPC due to the absence of a window size requirement (Glerean, Salmi et al. 2012, Omidvarnia, Pedersen et al. 2016, Pedersen, Omidvarnia et al. 2018, Zhou, Zhu et al. 2023) does not take into account the sensitivity of PS to bandwidth and frequency band selection. For instance, in our simulation analysis (Figure 3), bandwidths BW1 and BW2 perform poorly in Scenario 1, where PS was expected to outperform SWPC, highlighting PS’s sensitivity to bandwidth selection. The concept of the “right” bandwidth for PS remains unresolved, making it challenging to recommend PS over SWPC for time-resolved functional connectivity analysis especially when there is evidence that suggests the bandwidth of neural relevant information varies across subjects (Handwerker, Ollinger and D’Esposito 2004, Grady and Garrett 2014). Furthermore, although a 30-second window size for PS shows a high correlation with SWPC as demonstrated in Figure 4 (A), the clinical relevance provided by SWPC using a longer window size, such as 88 seconds, is not captured by PS (see supplementary material 5). This indicates that the similarities between the two methods do not inherently make one superior to the other.

Moreover, the concept of instantaneous phase is straightforward to interpret for monochromatic (single-frequency) signals, whereas the notion of a single-valued instantaneous frequency and phase is not meaningful for all types of signals (Boashash 1992, Picinbono 1997). This underscores the necessity of narrowband filtering when employing the Hilbert transform (Boashash 1992, Chavez, Besserve et al. 2006) for computing PS in fMRI studies. Therefore, selecting an appropriate narrow frequency band is essential to obtain a meaningful interpretation of phase synchronization from fMRI data. The recommendation to use PS (Glerean, Salmi et al. 2012, Omidvarnia, Pedersen et al. 2016, Pedersen, Omidvarnia et al. 2017, Pedersen, Omidvarnia et al. 2018) overlooks the critical parameter of frequency band selection. It suggests that PS is superior to SWPC because it does not require choosing a window size, yet it neglects the necessity of constricting the frequency band of the fMRI signal for meaningful PS analysis (Glerean, Salmi et al. 2012). Note that window size and shape selection in SWPC is in essence a filter design procedure (Faghiri, Iraji et al. 2022) and therefore is quite similar to the filter design step that is necessary for estimating PS using the Hilbert transform. Additionally, estimating instantaneous phase based on the analytic signal definition is possible for specific types of signals which makes it inapplicable to all types of signals (Picinbono 1997). Future work needs to investigate if fMRI signals match these assumptions.

Considering the limitations highlighted for the phase synchronization method, we propose an alternative perspective on SWPC and PS: neither method holds superiority over the other. We conclude that the fundamental differences between PS and SWPC necessitate that they be studied separately to uncover complementary insights into time-resolved functional network connectivity and the brain. Thus, we recommend further studies explore phase-amplitude trFNC measures to fully leverage their unique contributions.

A limitation of this study was our inability to quantify the exact reasons for the dissimilarities observed between the two methods in some cases. We propose several possible reasons, such as necessary frequency mismatches, the oscillatory properties of the fMRI data, and different connectivity properties, as demonstrated in our simulations. However, it remains unclear how each of these factors contributes to the observed differences and to what degree. We recommend that future research focuses on quantifying the effects of these factors on the observed differences to better understand their impact in trFNC analyses. We also encourage future research to incorporate both cosine and sine aspects of the phase difference to give a more accurate phase synchronization measure rather than focusing on the cosine-based phase methods.

## 7. CONCLUSION

In conclusion, our study underscores the complementary nature of SWPC and PS, each leveraging distinct signal properties—amplitude and phase, respectively—to illuminate brain network interactions. Integrating the strengths of both methods promises a more comprehensive understanding of time-resolved functional connectivity in fMRI research.

## 9. AUTHORS

**Sir-Lord Wiafe:** Conceptualization, Formal analysis, Methodology, Visualization, Writing – original draft. **Nana Asante:** Methodology, Writing –review & editing. **Vince Calhoun:** Conceptualization, Funding acquisition, Validation, Methodology, Resources, Supervision, Writing – review & editing. **Ashkan Faghiri:** Conceptualization, Methodology, Supervision, Validation, Writing –review & editing.

## 10. DATA & CODE AVAILABILITY STATEMENT

The codes for all our analyses in MATLAB language can be accessed through GitHub (https://github.com/Sirlord-Sen/studying_trFNC_via_communication_theory). The data was not collected by us and was provided in a deidentified manner. The IRB will not allow sharing of data or individual derivatives as a data reuse agreement was not signed by the subjects during the original acquisition.

## 11. ACKNOWLEGDEMENT

This work was supported by the National Institutes of Health (NIH) grant (R01MH123610), the National Science Foundation (NSF) grant #2112455.

