## supplementary material 1 for "Studying time-resolved functional connectivity via communication theory: on the complementary nature of phase synchronization and sliding window Pearson correlation"

*Proof of SWPC as an amplitude demodulation technique*

Given a message signal $m(t)$, carrier signal $c(t)$, and amplitude modulated signal $y(t)$, the relationship between $SWPC(c\left( t \right), y(t))$ and $m(t)$ given a window length ($\Delta$), $2L+1$, is shown below:

$$c\left( t \right)= A_{c}\cos2\pi f_{c}t, y\left( t \right)= \left( 1+ \frac{m\left( t \right)}{A_{c}} \right)c\left( t \right)$$

$$SWPC\left( y,c;t \right)= \frac{\sum_{\tau=t-L}^{t+L} \left( y_{\tau}- \bar{y_{\tau}} \right)\left( c_{\tau}- \bar{c_{\tau}} \right)}{\sqrt{\sum_{\tau=t-L}^{t-L-1} \left( y_{\tau}- \bar{y_{\tau}} \right)^{2}\left( c_{\tau}- \bar{c_{\tau}} \right)^{2}}}$$

$$SWPC\left( y,c;t \right)= \frac{\sum_{\tau=t-L}^{t+L} \left( \left( 1+ \frac{m_{\tau}}{A_{c}} \right)c_{\tau}- \bar{y_{\tau}} \right)\left( c_{\tau}- \bar{c_{\tau}} \right)}{\sqrt{\sum_{\tau=t-L}^{t+L} \left( \left( 1+ \frac{m_{\tau}}{A_{c}} \right)c_{\tau}- \bar{y_{\tau}} \right)^{2}\left( c_{\tau}- \bar{c_{\tau}} \right)^{2}}}$$

$$\bar{y_{\tau}}= \frac{1}{\Delta}\sum_{\tau=t-L}^{t+L} \left( 1+ \frac{m_{\tau}}{A_{c}} \right)A_{c}\cos2\pi f_{c}\tau$$

$$\bar{y_{\tau}}= \frac{A_{c}}{\Delta}\sum_{\tau=t-L}^{t+L} \left( 1+ \frac{m_{\tau}}{A_{c}} \right)\sum_{\tau=t-L}^{t+L} \cos2\pi f_{c}\tau$$

$$\bar{c_{\tau}}= \frac{A_{c}}{\Delta}\sum_{\tau=t-L}^{t+L} \cos2\pi f_{c}\tau$$

$$for large \Delta, \frac{1}{\Delta}\sum_{\tau=t-L}^{t+L} \cos2\pi f_{c}\tau\cong0$$

$$SWPC\left( y,c;t \right)= \frac{\sum_{\tau=t-L}^{t+L} \left( 1+ \frac{m_{\tau}}{A_{c}} \right){c_{\tau}}^{2}}{\sqrt{\sum_{\tau=t-L}^{t+L} \left( 1+ \frac{m_{\tau}}{A_{c}} \right)^{2}{c_{\tau}}^{4}}}$$

$$SWPC\left( y,c;t \right)= \frac{\sum_{\tau=t-L}^{t+L} {c_{\tau}}^{2}\sum_{\tau=t-L}^{t+L} \left( 1+ \frac{m_{\tau}}{A_{c}} \right)}{\sqrt{\sum_{\tau=t-L}^{t+L} {c_{\tau}}^{4}\sum_{\tau=t-L}^{t+L} \left( 1+ \frac{m_{\tau}}{A_{c}} \right)^{2}}}$$

$$\sum_{\tau=t-L}^{t+L} {c_{\tau}}^{2}= \sum_{\tau=t-L}^{t+L} {A_{c}}^{2}\cos^{2} 2\pi f_{c}\tau= {A_{c}}^{2}\Delta\sum_{\tau=t-L}^{t+L} \cos^{2} 2\pi f_{c}\tau\propto{{A_{c}}^{2}\Delta}^{2}$$

$$\sum_{\tau=t-L}^{t+L} {c_{\tau}}^{4}= \sum_{\tau=t-L}^{t+L} {A_{c}}^{4}\cos^{4} 2\pi f_{c}\tau={A_{c}}^{4}\Delta\sum_{\tau=t-L}^{t+L} \cos^{4} 2\pi f_{c}j \propto{A_{c}}^{4}\Delta^{2}$$

$$SWPC\left( y,c;t \right)\propto\frac{{A_{c}}^{2}\Delta^{2}\sum_{\tau=t-L}^{t+L} \left( 1+ \frac{m_{\tau}}{A_{c}} \right)}{\sqrt{{A_{c}}^{4}\Delta^{2}\sum_{\tau=t-L}^{t+L} \left( 1+ \frac{m_{\tau}}{A_{c}} \right)^{2}}}$$

$$SWPC\left( y,c;t \right)\propto\frac{{A_{c}}^{2}\Delta^{2}\sum_{\tau=t-L}^{t+L} \left( 1+ \frac{m_{\tau}}{A_{c}} \right)}{{A_{c}}^{2}\Delta\sqrt{\sum_{\tau=t-L}^{t+L} \left( 1+ \frac{m_{\tau}}{A_{c}} \right)^{2}}}$$

$$SWPC\left( y,c;t \right)\propto\Delta\frac{\sum_{\tau=t-L}^{t+L} \left( 1+ \frac{m_{\tau}}{A_{c}} \right)}{\sqrt{\sum_{\tau=t-L}^{t+L} \left( 1+ \frac{m_{\tau}}{A_{c}} \right)^{2}}}$$
