## supplementary material 2 for "Studying time-resolved functional connectivity via communication theory: on the complementary nature of phase synchronization and sliding window Pearson correlation"

*Proof of phase modulation being a non-linear modulation scheme using the principle of superposition*

Phase modulation (PM), unlike AM, modulates the phase of the carrier signal with the amplitude of the message signal. Mathematically, this can be expressed as:

$$y\left( t \right)=A_{c}\cos\left( 2\pi f_{c}t+k_{p}m(t) \right)$$

Where $k_{p}$ is the phase sensitivity of the modulator, a constant that determines how much the phase of the carrier will be affected (Haykin and Moher 1989). Assuming $k_{p}$ to be 1 for simplicity, the phase modulated signal becomes:

$$y\left( t \right)=A_{c}\cos\left( 2\pi f_{c}t+m(t) \right)$$

Given a message signal $m\left( t \right)$ which is composed of two components $m_{1}\left( t \right)$ and $m_{2}\left( t \right)$ as follows:

$$m\left( t \right)= m_{1}\left( t \right)+ m_{2}\left( t \right)$$

Applying the superposition theorem, each of the signals will be phase modulated and if the superposition theorem is obeyed, the phase modulated signal of $m\left( t \right)$, $y\left( t \right)$, should be equal to the phase modulated signal of $m_{1}\left( t \right)$, $y_{1}\left( t \right)$ and phase modulated signal of $m_{2}\left( t \right)$, $y_{2}\left( t \right)$.

The phase modulation of $y\left( t \right)$ as given below ($k_{p}=1$):

$$y\left( t \right)= A_{c}\cos\left( 2\pi f_{c}t+m_{1}\left( t \right)+ m_{2}\left( t \right) \right)$$

$$y\left( t \right)= A_{c}\cos\left( 2\pi f_{c}t+ m\left( t \right) \right)$$

$$y\left( t \right)= A_{c}\left[ \cos\left( 2\pi f_{c}t \right)\cos\left( m\left( t \right) \right)- \cos\left( 2\pi f_{c}t \right)\cos\left( m\left( t \right) \right) \right]$$

The phase modulation of $y_{1}\left( t \right)+ y_{2}\left( t \right)$ is given below:

$$y_{1}\left( t \right)= A_{c}\cos\left( 2\pi f_{c}t+m_{1}\left( t \right) \right), y_{2}\left( t \right)= A_{c}\cos\left( 2\pi f_{c}t+m_{2}\left( t \right) \right)$$

$$y_{1}\left( t \right)+ y_{2}\left( t \right)= A_{c}\cos\left( 2\pi f_{c}t+m_{1}\left( t \right) \right)+ A_{c}\cos\left( 2\pi f_{c}t+m_{2}\left( t \right) \right)$$

$$y_{1}\left( t \right)+ y_{2}\left( t \right)=2A_{c}\left[ \cos\frac{4\pi f_{c}t+ m\left( t \right)}{2}\cos\frac{m_{1}\left( t \right)- m_{2}\left( t \right)}{2} \right]$$

From the above expressions, it is easily seen that though $m\left( t \right)= m_{1}\left( t \right)+ m_{2}\left( t \right)$, the superposition theorem is not obeyed under phase modulation emphasizing its nonlinearity since:

$$y_{1}\left( t \right)+ y_{2}\left( t \right) \neq y\left( t \right)$$
