## supplementary material 3 for "Studying time-resolved functional connectivity via communication theory: on the complementary nature of phase synchronization and sliding window Pearson correlation"

*Estimated probability density functions of SWPC and PS across all three cases.*

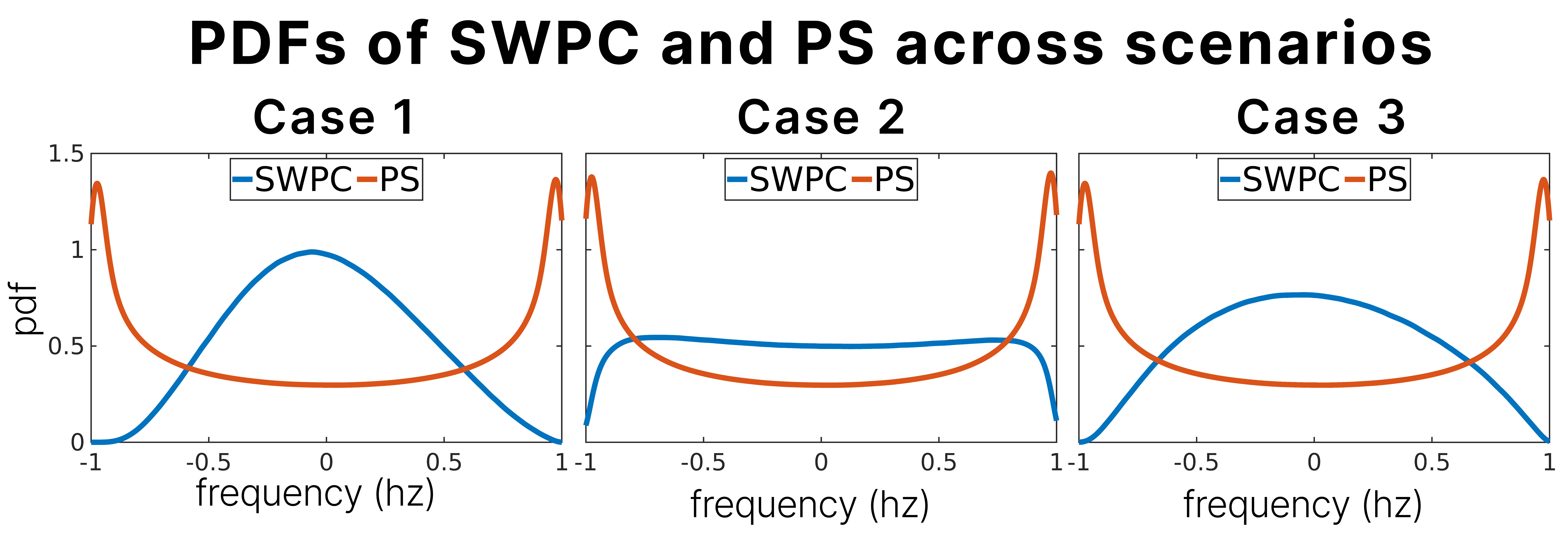

The estimated probability density function (PDF) of SWPC and PS estimations is shown above. Given that the PS distributions are not normal in all cases, using Spearman correlation will provide more reliable similarity coefficients between SWPC and PS.
