## supplementary material 4 for "Studying time-resolved functional connectivity via communication theory: on the complementary nature of phase synchronization and sliding window Pearson correlation"

*Elbow plots for SWPC and PS across all three cases.*

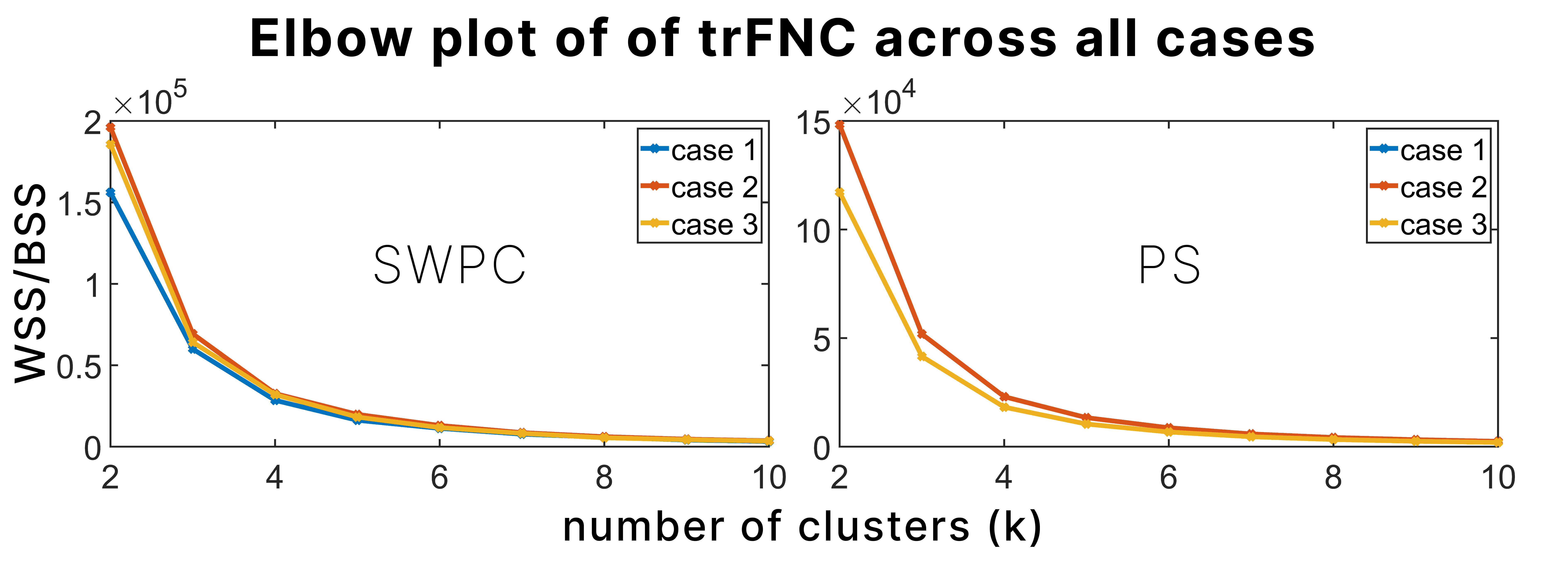

The elbow plot from k-means clustering, using clusters ranging from 2 to 10, and the ratio of the within-cluster sum of squared distances (WSS) to the between-cluster sum of squared distances (BSS) for SWPC and PS across all cases, indicates that the optimal number of clusters is 4.
