## supplementary material 5 for "Studying time-resolved functional connectivity via communication theory: on the complementary nature of phase synchronization and sliding window Pearson correlation"

*trFNC and group analysis of SWPC and PS across all three cases.*

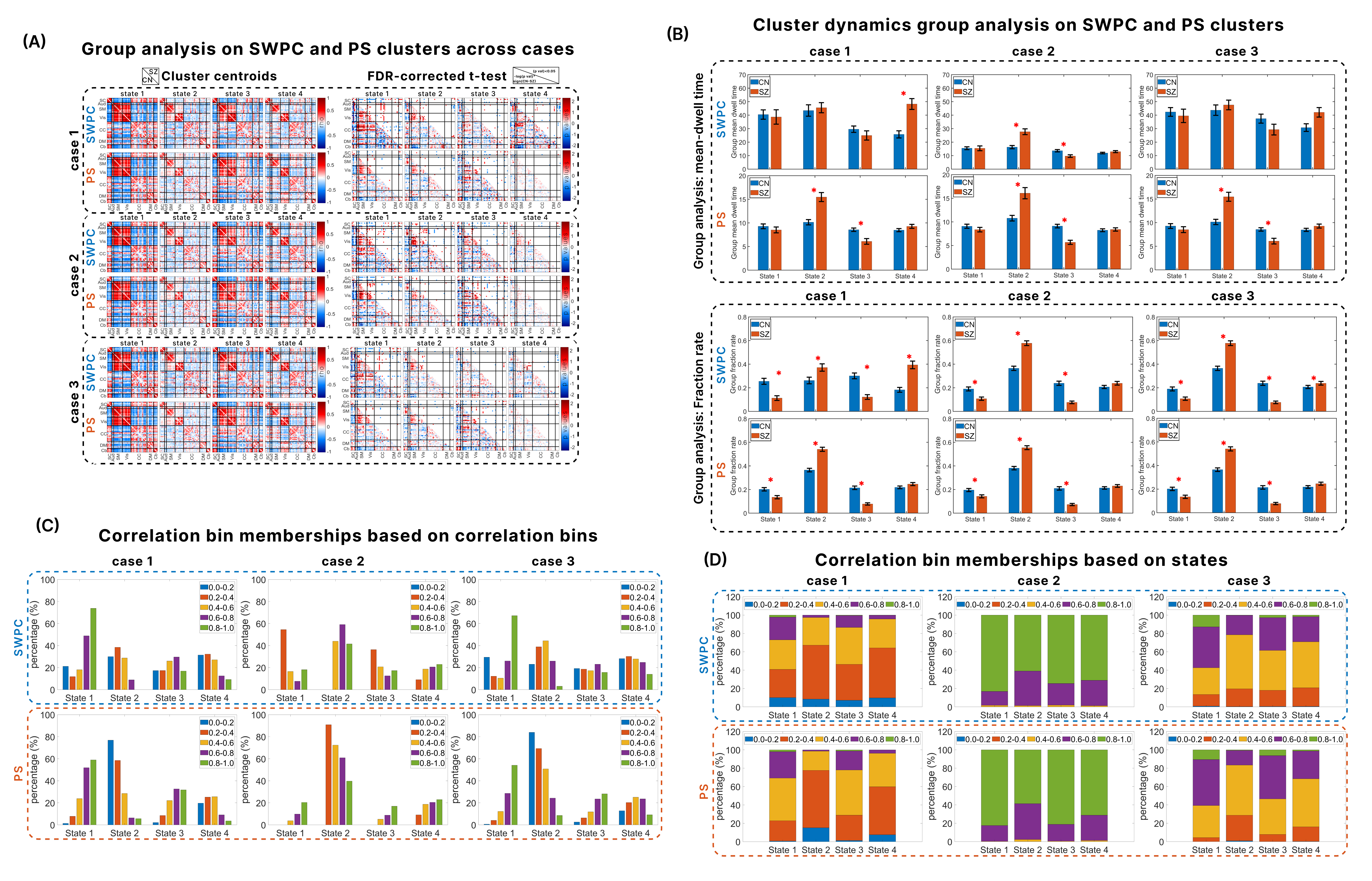

After computing SWPC and PS across all cases, k-means clustering with 4 clusters was applied independently to each method to obtain 4 clusters/states. These states were categorized into corresponding states representing schizophrenia and control groups. A two-sample t-test was then performed between the states from the schizophrenia group and the control group. The clusters/states are presented on the left side of (A), while the false discovery rate (FDR) corrected p-values from the t-test are displayed on the right side of (A).

(B) illustrates the mean-dwell time and fraction rate of SWPC and PS across all cases between schizophrenia and controls. We divided the time points of SWPC and PS into bins reflecting the level of correlation between the two methods using Spearman correlation. The bins are 0-0.2, 0.2-0.4, 0.4-0.6, 0.6-0.8, and 0.8-1. For each case, we computed the constituency of each correlation bin within the cluster as a fraction of the correlation bin, as shown in (C), while (D) shows the constituency of each correlation bin within the cluster as a fraction of the time points in that state.

A key takeaway is that in case 2, where the correlation between SWPC and PS is the most similar (as noted in the main manuscript), the results for mean dwell time (MDT) and fraction rate (FR) are also similar. This might suggest that the two methods are comparable when they yield similar results. However, this similarity causes the significance of MDT in SWPC for case 1 to be lost, indicating that a strong correlation between the two methods is not necessarily a reliable way to evaluate them. The complementary results observed in cases 1 and 3 further support this point.
