## supplementary material 6 for "Studying time-resolved functional connectivity via communication theory: on the complementary nature of phase synchronization and sliding window Pearson correlation"

*Simulation results using root mean squared error as an evaluation metric.*

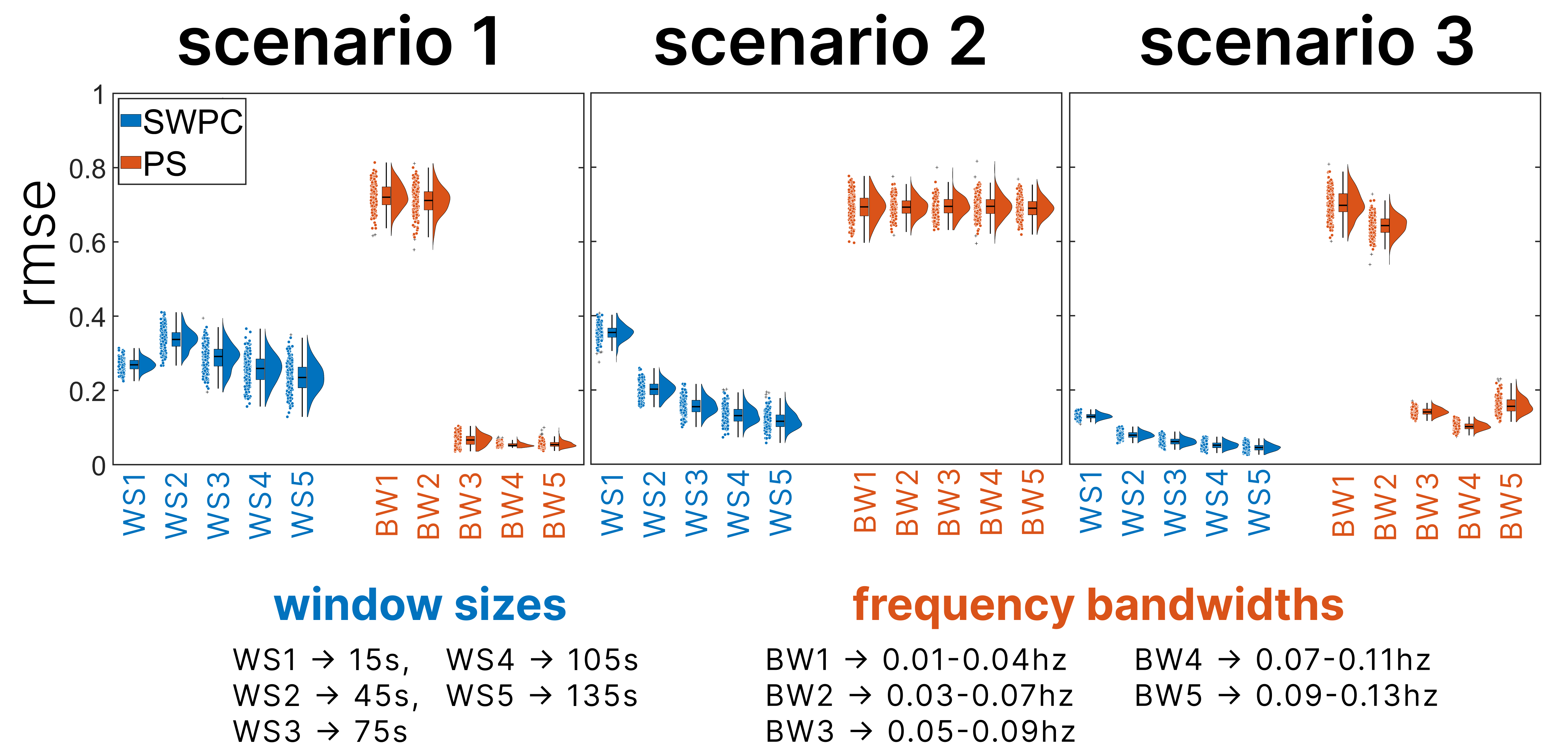

We created three scenarios to evaluate the performance of SWPC and PS with different time-resolved connectivity encoded between a pair of signals. Scenario 1 involves low-magnitude high-frequency connectivity encoded in the phase of the signal pairs. Scenario 2 involves high-amplitude low-frequency connectivity encoded in the amplitude of the signal pairs. Scenario 3 involves mid-magnitude mid-frequency connectivity encoded in both the phase and amplitude of the signal pairs.

As shown in the main manuscript, based on correlation, PS performs better in scenario 1, SWPC performs better in scenario 2, and both methods perform well in scenario 3. These results are supported by the root mean squared error (RMSE) of the estimated connectivity compared to the ground truth across all three scenarios.
